# Medulloblastoma-Associated KBTBD4 Mutations Disrupt PP2A-A Orphan Quality Control

**DOI:** 10.64898/2026.03.02.709011

**Authors:** Regina Baur, Laura A. Schneider, Gajanan Sathe, Thomas Lunardi, Jonathan Schneider, Anna-Sophia Krebs, Juan C. Silva, Diane L. Haakonsen, Sebastian M. Waszak, Joachim Lingner, Alessio Ciulli, Michael Rape, Nicolas H. Thomä

**Affiliations:** School of Life Sciences, École Polytechnique Fédérale de Lausanne (EPFL), Lausanne, Switzerland; Department of Molecular and Cell Biology, University of California at Berkeley, Berkeley, CA, USA; Howard Hughes Medical Institute, University of California at Berkeley, Berkeley, CA, USA; Faculty of Biosciences, Heidelberg University, Heidelberg, Germany; Centre for Targeted Protein Degradation, School of Life Sciences, University of Dundee, 1 James Lindsay Place, Dundee DD1 5JJ, United Kingdom; Molecular Biology Institute, UCLA, Los Angeles, California, USA; Lunenfeld-Tanenbaum Research Institute, Sinai Health System, Toronto, Ontario, Canada; Friedrich Miescher Institute for Biomedical Research, Basel, Switzerland

## Abstract

Medulloblastoma, the most common malignant pediatric brain tumor, arises from developmental aberrations of cerebellar precursor cells. The CUL3-RING ubiquitin ligase adaptor KBTBD4 is recurrently mutated in medulloblastoma subgroups 3 and 4. While *KBTBD4* mutations confer a gain-of-function phenotype leading to aberrant degradation of transcriptional repressors, endogenous targets of this E3 ligase remain unknown. Here, we identify the PP2A-A scaffolding subunit of the PP2A phosphatase as a CRL3^KBTBD4^ substrate. Using a combination of proteomics, cell biology, biochemical reconstitution, and cryo-EM structural analyses, we show that CRL3^KBTBD4^ mediates orphan quality control by targeting free PP2A-A for degradation to safeguard phosphatase activity. Loss of *KBTBD4* or its mutation in medulloblastoma cause PP2A-A accumulation, hence affecting phospho-dependent signaling pathways in cancer development. Disease mutations in KBTBD4 thus elicit a dual phenotype: gain-of-function degradation of transcriptional repressors combined with loss of PP2A quality control, which dysregulates multiple signaling events implicated in cancer, including telomere length regulation.

## Introduction

Pediatric brain cancers have surpassed leukemia as the leading cause of cancer-related mortality in children (Curtin et al. 2016). Among embryonal tumors of the central nervous system, medulloblastomas (MBs) are the most prevalent. MBs are subdivided into four molecular subgroups, WNT, SHH, Group 3, and Group 4, that show unique genetic and clinical features (Northcott et al. 2011; Northcott, Shih, et al. 2012; Northcott, Dubuc, et al. 2012; Northcott et al. 2014). The WNT and SHH subgroups are primarily driven by mutations in *CTNNB1* and *PTCH1*, respectively, whereas Group 3 and Group 4 MBs harbor alterations in transcriptional regulators such as *GFI1/GFI1B*, *PRDM6*, *MYC/MYCN*, as well as in epigenetic modifiers including *KDM6A* (Sharma et al. 2019). Recurrent mutations in *KBTBD4*, a substrate adaptor of a CUL3 RING E3 ubiquitin ligase, have also been identified as drivers in Group 3 and Group 4 medulloblastomas (Northcott et al. 2017), as well as in pineal parenchymal tumors (Lee et al. 2019). Although patients with Group 3 and Group 4 MB tumors have poor clinical outcomes, the molecular mechanisms of oncogenic events driven by mutations in *KBTBD4* remain incompletely understood.

Recent studies found that KBTBD4 insertion and deletion (indel) mutations confer a gain-of-function phenotype that results in the aberrant ubiquitin-dependent degradation of the CoREST complex (Chen et al. 2022). CoREST is a central epigenetic corepressor that prevents neuronal differentiation and plays a central role in several tumors with neurodevelopmental origin, including medulloblastoma, by maintaining a stem-cell like phenotype of cancer cells (Andrés et al. 1999; Park et al. 2022; Ismail et al. 2025). A similar gain-of-function effect can be induced pharmacologically by the small-molecule degrader UM171, which, in conjunction with wild-type KBTBD4, promotes CoREST degradation (Chagraoui et al. 2021). Both UM171 treatment and MB-associated mutations enhance KBTBD4 binding to HDAC1, thereby stimulating ubiquitylation and proteasomal degradation of CoREST subunits (Yeo et al. 2025; Xie et al. 2025; Chen et al. 2025).

Most gain-of-function mutations in cancer disrupt physiological regulation of a signaling pathway to increase its natural output. As an example, mutations in β-catenin impair recognition by its native E3 ligase SCF^βTrCP^, thereby prolonging activity of this transcription factor and eliciting a gene expression program that is independent of WNT signals (Liu et al. 1999). Surprisingly, neither HDAC1 nor other CoREST subunits appear to be regulated by wild-type CRL3^KBTBD4^, and physiological substrates of a KBTBD4-dependent E3 ligase have not been discovered. Whether mutations in *KBTBD4* disrupt endogenous signaling pathways remains unknown, leaving the impact of *KBTBD4* in MB formation incompletely understood.

Here, we show that KBTBD4 is a core component of an orphan quality control pathway that safeguards subunit stoichiometry of the PP2A phosphatase complex. PP2A is a ubiquitously expressed tumor suppressor itself and critical for most essential cellular processes such as mitosis and cell division, signal transduction and differentiation (Brautigan and Shenolikar 2018; Kauko and Westermarck 2018). CRL3^KBTBD4^ targets the central scaffolding subunit of PP2A, PP2A-A, yet only when PP2A-A is not incorporated into the heterotrimeric phosphatase complex. This quality control pathway is disrupted by either MB mutations in *KBTBD4* or mutations in *PPP2R1A* that have been found in uterine cancers (Taylor et al. 2019). Thus, oncogenic mutations in *KBTBD4* simultaneously elicit gain-of-function degradation of CoREST and loss of PP2A quality control, a complex outcome of a single mutation that effectively reshapes multiple critical signaling networks in medulloblastoma.

## Results

### CRL3^KBTBD4^ targets the PP2A phosphatase

To identify physiological targets of CRL3^KBTBD4^, we purified transiently expressed KBTBD4 from HEK293T cells and identified its interactome by mass spectrometry. Inhibiting the Cullin system with the neddylation inhibitor MLN4924, and the proteasome system with Carfilzomib (CFZ), we observed increased binding of the scaffolding subunit PP2A-A of the phosphatase PP2A to KBTBD4 (**Fig. 1A**). In parallel, we generated Δ*KBTBD4* cell lines from parental HEK293T cells (**Fig. S1A**) and used mass spectrometry to compare total protein levels in Δ*KBTBD4* cells to those of their wild-type (WT) counterparts. Here, we noticed stabilization of PP2A-A in two Δ*KBTBD4* clonal cell lines (**Fig. 1B**). We confirmed elevated PP2A-A levels in Δ*KBTBD4* cells by western blot, whereas levels of the catalytic subunit of PP2A (PP2A-C) or a substrate-specificity factor (PP2A-B) remained unperturbed (**Fig. 1C**). As PP2A is an abundant and established tumor suppressor with a wide repertoire of phospho-substrates to globally antagonize mitogenic kinase signaling (Brautigan and Shenolikar 2018; Kauko and Westermarck 2018; Brewer et al. 2024), we focused on the functional and mechanistic link between CRL3^KBTBD4^ and phospho signaling governed by the PP2A phosphatase.

**Figure 1:**
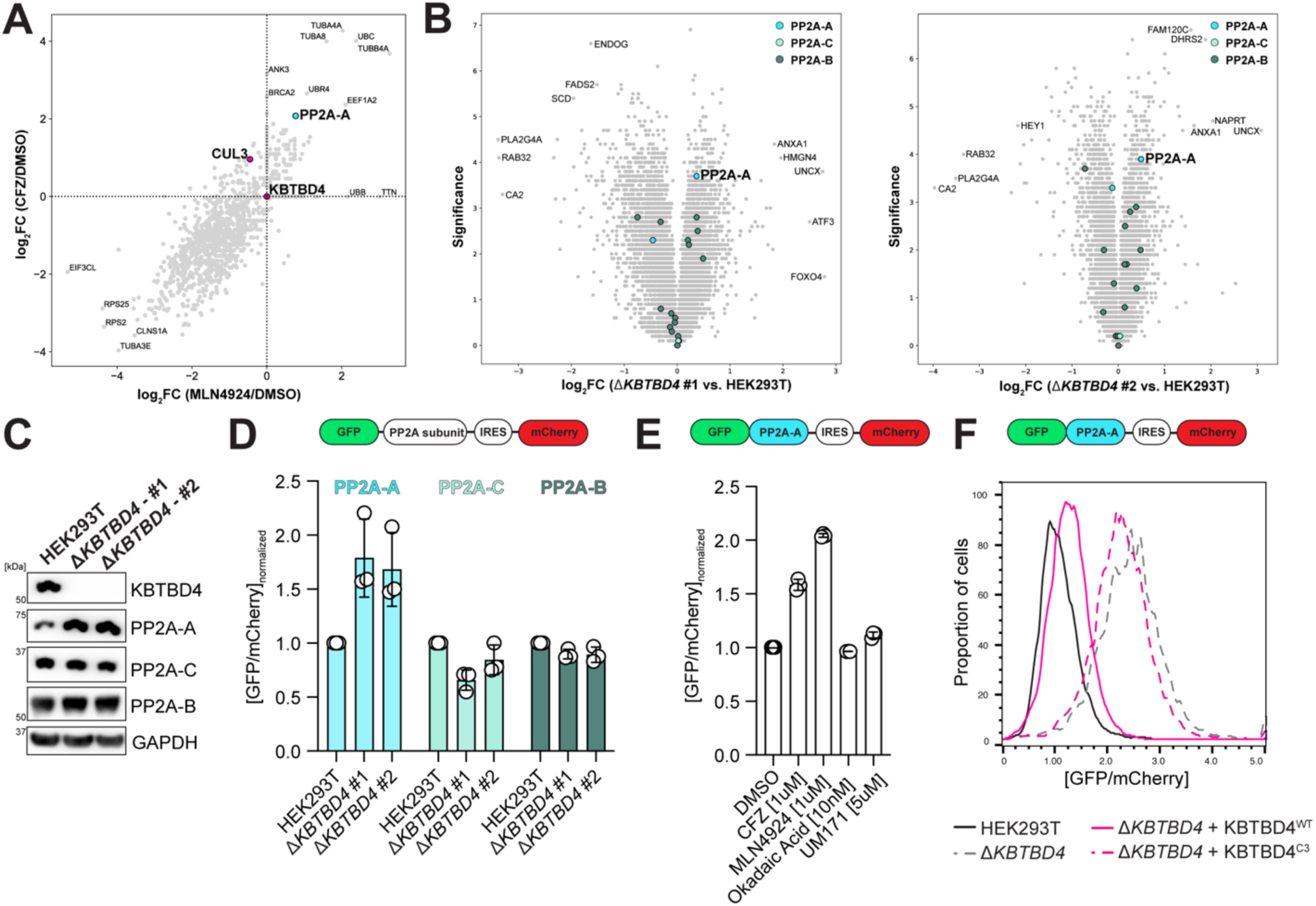
PP2A-A is an interactor and substrate of CRL3^KBTBD4^. **A.** Transiently expressed KBTBD4^3×Flag^ was purified from DMSO-, MLN4924- and CFZ-treated HEK293T cells and interactors were identified by mass spectrometry. Spectral counts were normalized to bait count, and log_2_ fold changes of MLN4924 and CFZ treatment relative to control were plotted. **B.** Whole cell extracts from Δ*KBTBD4* cells #1 and #2 were analyzed in triplicates using TMT-based proteomics to show quantitative changes of protein levels. The log_2_ fold change of protein abundance is plotted against the significance (-log_10_(p-value)) as determined with a two-tailed t-test. **C.** Whole cell extracts from HEK293T and Δ*KBTBD4* cell lines #1 and #2 were analyzed by western blotting with the indicated antibodies. Representative result from n=3 independent experiments. **D.** Stability of the PP2A complex subunits PP2A-Aα, PP2A-Cα, and PP2A-B55α was measured in HEK293T and Δ*KBTBD4* cell lines by transient expression of the dual fluorescence reporter in flow cytometry. Data was measured in triplicates, and the median fluorescence signal ratio of GFP over mCherry was normalized to HEK293T control. **E.** Stability of PP2A-A was measured in HEK293T cells with the dual fluorescence reporter assay in flow cytometry after addition of the indicated drugs. Data was measured in triplicates, and the median fluorescence signal ratio of GFP over mCherry was normalized to DMSO. **F.** PP2A-Aα stability was monitored by flow cytometry through transient expression of the dual fluorescence reporter in HEK293T cells and Δ*KBTBD4* #1 with co-expression of wild-type KBTBD4^WT^ or the CUL3-binding deficient KBTBD4^C3^ construct. Similar results in n=3 independent samples.

To determine whether CRL3^KBTBD4^ directly regulates PP2A turnover, we made use of a protein stability reporter (Yen and Elledge 2008; Yen et al. 2008; Manford et al. 2020; Haakonsen et al. 2024), and found that deletion of *KBTBD4* resulted in robust stabilization of the scaffolding subunit PP2A-A (**Fig. 1D**). The PP2A-B and PP2A-C subunits were not strongly affected (**Fig. 1D**). The effects of *KBTBD4* deletion on PP2A-A stability were comparable to treatment of cells with CFZ or MLN4924, while PP2A-A stability was not affected by UM171 or PP2A inhibition by okadaic acid (**Fig. 1E**). Importantly, PP2A-A turnover could be restored in Δ*KBTBD4* cells by re-expression of wild-type KBTBD4, whereas a CUL3-binding deficient variant (KBTBD4^C3^) did not rescue PP2A-A levels (**Fig. 1F**). These results demonstrate that the E3 ligase activity of CRL3^KBTBD4^ controls stability and abundance of PP2A-A.

### CRL3^KBTBD4^ ubiquitylates unassembled PP2A-A

The PP2A phosphatase is a heterotrimeric complex: the scaffold PP2A-A serves as a docking platform for the catalytic subunit PP2A-C and the substrate adaptor PP2A-B to recruit and dephosphorylate specific substrates (Xu et al. 2006; Cho and Xu 2007). With two PP2A-A, 2 PP2A-C, and ∼16 PP2A-B isoforms, PP2A comprises up to ∼100 possible heterotrimeric phosphatase complexes (Sangodkar et al. 2016). PP2A biogenesis is tightly regulated and requires assembly of a PP2A-AC core, followed by incorporation of regulatory PP2A-B subunits (Haanen et al. 2022). The rogue activity of free PP2A-C is prevented by sequestration through endogenous inhibitors (Kong et al. 2009; Kauko and Westermarck 2018; Haanen et al. 2022; Wachter et al. 2024).

To assess how CRL3^KBTBD4^ impacts PP2A, we investigated the interaction of KBTBD4 with representative members of hetero-trimeric PP2A complex. We choose the most abundant PP2A-A and PP2A-C isoforms, PP2A-Aα and -Cα, respectively, and purified them either individually or as dimeric PP2A-AC complex. We also purified PP2A-AC bound to the substrate adaptor B55α as a representative of a heterotrimeric PP2A complex (**Fig. 2A, S1B, C**). KBTBD4 associated with free recombinant PP2A-A, whereas no interaction was detected between KBTBD4 and PP2A-B or PP2A-C, respectively (**Fig. 2A; Fig. S1B, C**). In the presence of subcomplexes, KBTBD4 engaged free excess PP2A-A, but did neither associate with excess PP2A-C or PP2A-B subunits, nor the formed PP2A-AC and PP2A-ACB complexes (**Fig. 2A; Fig. S1B, C**).

**Figure 2:**
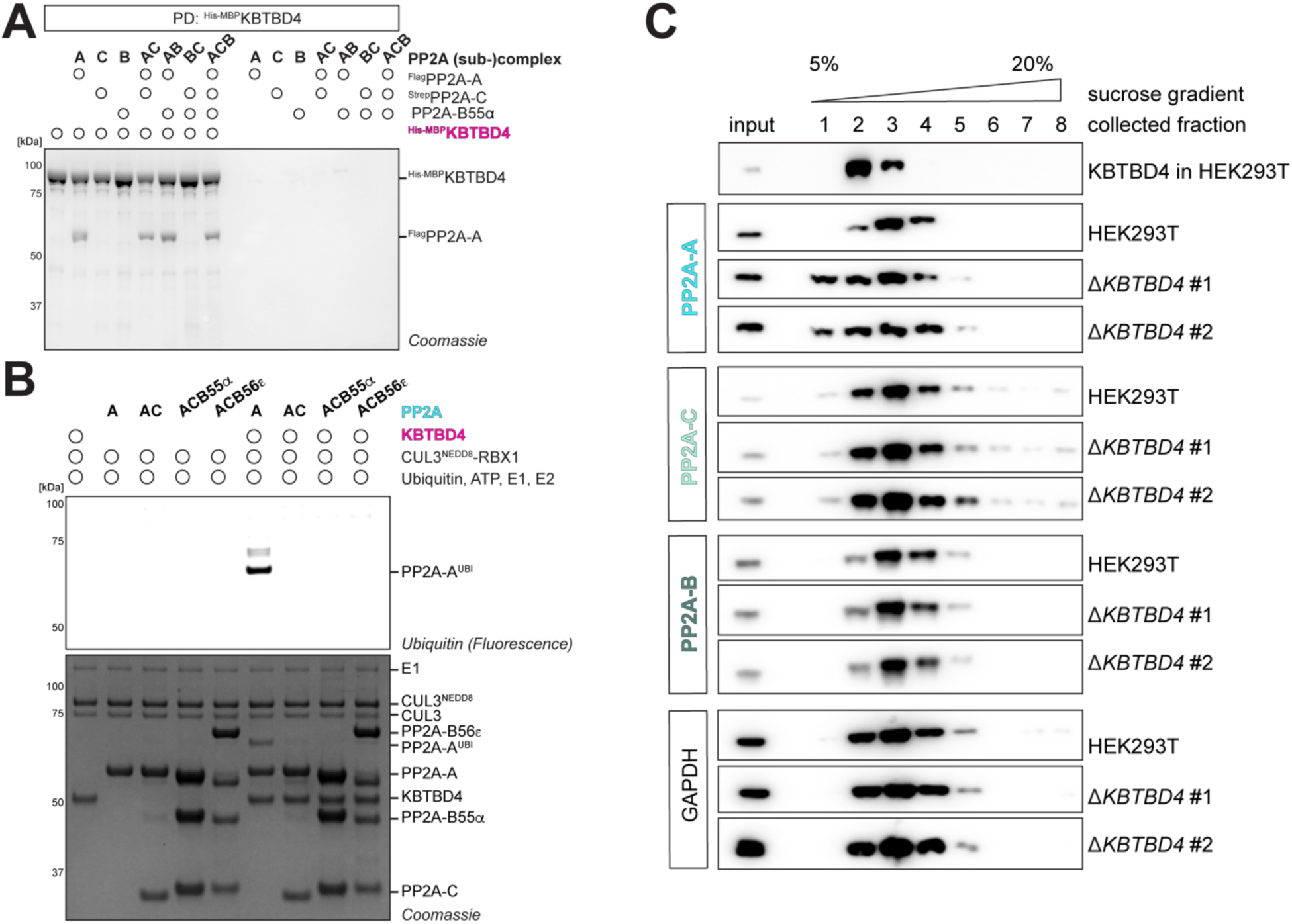
CRL3^KBTBD4^ ubiquitylates free PP2A-A. **A.** PP2A subunits PP2A-^3×Flag^Aα/^Strep^Cα/B55α were expressed as individual proteins, in PP2A subcomplexes PP2A-AC/AB/CB, and as the heterotrimeric PP2A-ACB holoenzyme in insect cells. PP2A extracts were mixed as indicated with ^6×His-MBP^KBTBD4 extract and affinity purified to test for binding of KBTBD4 to PP2A subunits and subcomplexes. **B.** Purified PP2A subunits and (sub-)complexes were ubiquitylated *in vitro* with purified KBTBD4, CUL3^NEDD8^-RBX1, UBE1, UBE2D3 and fluorescently labeled ubiquitin. Representative result of n=2 independent experiments. **C.** Cellular extracts from HEK293T cells and Δ*KBTBD4* cell lines #1and #2 were separated over a 5-20% sucrose gradient by ultracentrifugation. Fractions were probed for indicated proteins by western blot. Representative result of n=2 experiments.

To test whether PP2A-A binding translates into ubiquitylation by CRL3^KBTBD4^, we reconstituted the ubiquitylation cascade using purified proteins *in vitro*. We observed robust ubiquitylation of PP2A-A dependent on CRL3^KBTBD4^, but only if PP2A-A was present in isolation (**Fig. 2B**). CRL3^KBTBD4^ neither ubiquitylated the PP2A-AC core or heterotrimeric PP2A complexes (PP2A-ACB, **Fig. 2B**). As frequently seen with CUL3 E3 ligases (Werner et al. 2018; Rodríguez-Pérez et al. 2021; Akopian et al. 2022), CRL3^KBTBD4^ decorated PP2A-A with one or two ubiquitin moieties, and degradation may require further modification through chain-extending E3 ligases, such as the recently reported CUL3 partner TRIP12 (Ingersoll et al. 2025; Maiwald et al. 2025). By combining *in vitro* ubiquitylation with mass spectrometry, we found that CRL3^KBTBD4^ modified K34, K107, K188, K255, and K272 in PP2A-A (**Fig. S2A**). Importantly, while these Lys residues are readily accessible on the surface of PP2A-A, they are largely inaccessible within the heterotrimeric PP2A phosphatase (**Fig. S2B**). Sucrose gradient centrifugations of lysates of Δ*KBTBD4* compared to WT cells corroborated our findings (**Fig. 2C**). In WT cells, PP2A-A showed a sharp peak accompanied by PP2A-C and PP2A-B subunits corresponding to the PP2A holo-enzyme (**Fig. 2C**), while Δ*KBTBD4* cells had excess PP2A-A in lower molecular weight fractions that was not incorporated into heterotrimeric phosphatase complexes (**Fig. 2C**), indicating free excess PP2A-A.

Together, our results demonstrate that CRL3^KBTBD4^ preferentially binds and ubiquitylates unassembled PP2A-A. This behavior classifies CRL3^KBTBD4^ as an orphan quality control enzyme for the major architectural subunit of PP2A, specifically targeting orphan PP2A-A that has not been incorporated into PP2A complexes. This observation raised the possibility that orphan quality control through CRL3^KBTBD4^-dependent ubiquitylation of PP2A-A is required for optimal PP2A function and cellular phospho-homeostasis.

### Structural basis for protein complex quality control by CRL3^KBTBD4^

To visualize how CRL3^KBTBD4^ recognizes PP2A-A, we solved the cryo-EM structure of KBTBD4 bound to its substrate. Similar to most BTB proteins (Mena et al. 2018), mass photometry analysis showed that KBTBD4 primarily forms a dimer, while PP2A-A by itself exists as a monomer (**Fig. S2C**). Co-incubation of equimolar KBTBD4 and PP2A-A gave an observed mass of 256kDa, which was most consistent with the formation of a tetrameric complex composed of two molecules each of KBTBD4 and PP2A-A (**Fig. S2C**). As expected from our previous binding studies, neither the PP2A-AC core nor the complete PP2A-ACB phosphatase complex interacted with dimeric KBTBD4 in mass photometry, underscoring that KBTBD4 selectively recognizes PP2A-A molecules that have not been incorporated into PP2A complexes (**Fig. S2D**).

We subjected KBTBD4-PP2A-A complexes to mild BS3 crosslinking and obtained a cryo-EM structure with an overall resolution of 2.6 Å. In accordance with the mass photometry of the native complex, we observed a KBTBD4 dimer that engages two PP2A-A monomers (**Fig. 3A, B**). Each PP2A-A molecule interacts extensively with the Kelch domains of both KBTBD4 subunits along a C2 symmetry axis. Conversely, the two PP2A-A subunits within the tetrameric complex do not contact each other. We also observed a subpopulation of KBTBD4 engaging only a single PP2A-A molecule, which is sufficient to shift apo KBTBD4 to its substrate-binding conformation (**Fig. S2E**). The interaction of KBTBD4 with PP2A-A is centered on two main surfaces. One surface is formed by three top loops 2b-c, 3b-c, and 4b-c of the β-propellor repeats 2-4 of the Kelch domain (Kelch repeat; KR) of KBTBD4 with HEAT repeats 11-15 of one PP2A-A molecule; this interaction surface includes the residues within KBTBD4 loop 2b-c that are mutated in medulloblastoma (**Fig. 3C, D**). The second interaction patch is centered on the backside of the Kelch domain of KBTBD4, with the β-propellor repeats 1-3 engaging HEAT repeats 2-3 and 5-9 of the second PP2A-A molecule (**Fig. 3E, F**). Validating this structure, mutations of KBTBD4 residues at each interface with PP2A-A abrogated substrate recognition in immunoprecipitation experiments (**Fig. 3G**).

**Figure 3:**
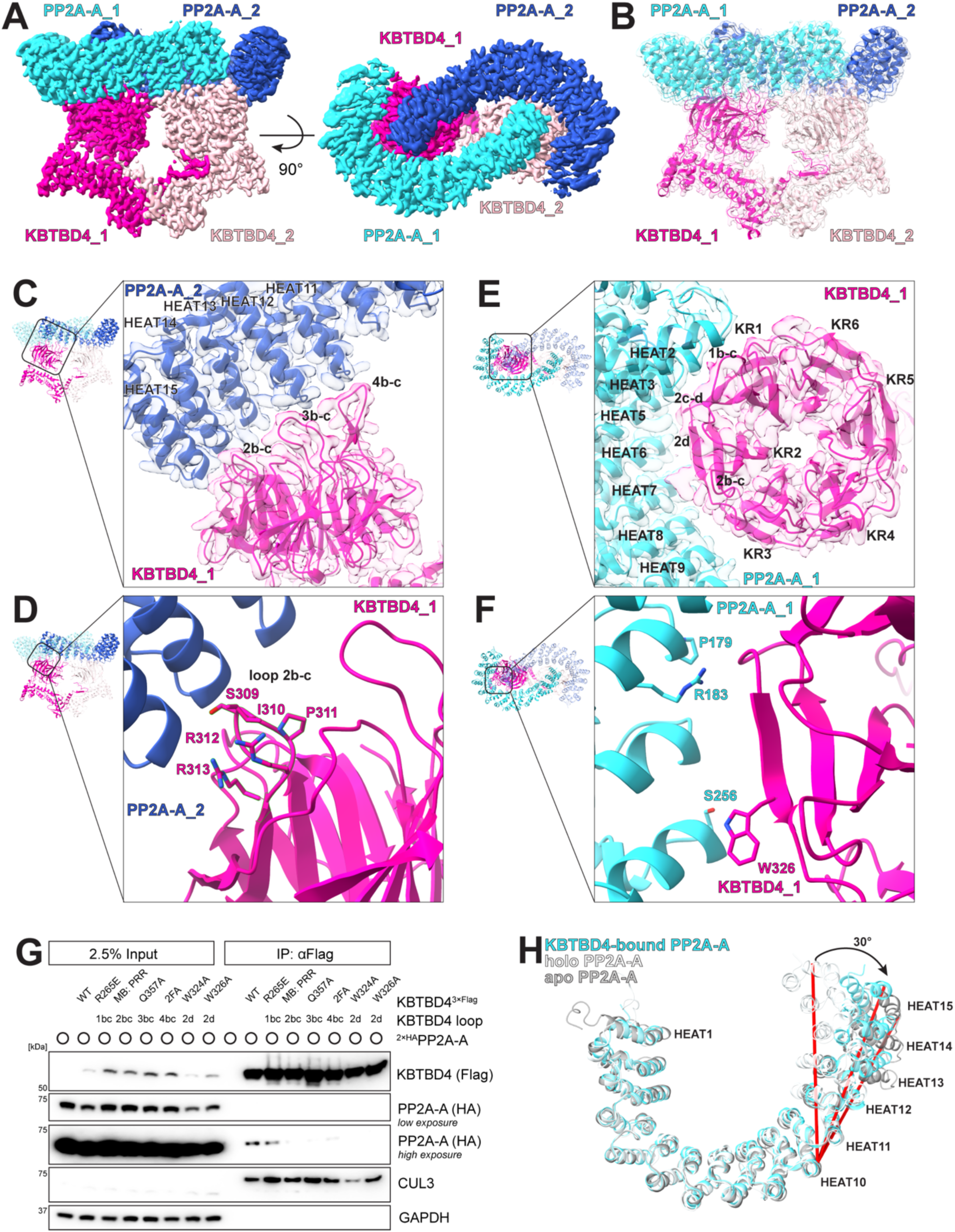
Dimeric KBTBD4 binds to two PP2A-A subunits. **A.** Cryo-EM map of a 2:2 complex of KBTBD4:PP2A-A in two orthogonal orientations at an overall resolution of 2.6 Å. KBTBD4 moieties in pink and rose, PP2A moieties in cyan and blue (EMDB: EMD-56545). **B.** Atomic model of the KBTBD4:PP2A-A complex derived from cryo-EM map in A (PDB: 28JO). **C.** Close up view of the top loops 2b-c, 3b-c, and 4b-c of the Kelch domain of KBTBD4_1 engaging the C-terminal HEAT repeats 11-15 of PP2A-A_2. **D.** Zoom into interface composed of loop 2b-c of KBTBD4_1, harboring MB hotspot mutations, and PP2A-A HEAT repeats 14 and 15. **E.** Close up view of the backside of the KBTBD4_1 Kelch domain engaging HEAT repeats 2-3 and 5-9 of PP2A-A_1 with its Kelch repeats (KR) 1-3. **F.** Close up view of KBTBD4_1:PP2A-A_1 interface containing PP2A-A HEAT repeat 5 with common cancer mutants P179 and R183. **G.** Co-immunoprecipitation of transiently expressed ^2×HA^PP2A-A and KBTBD4^3×Flag^ mutants to test effects of interaction surfaces as determined by cryo-EM. **H.** Comparison of the structure of monomeric apo PP2A-A (grey, PDB: 1B3U) to its conformations when bound to KBTBD4 (cyan, PDB: 28JO), or when incorporated into a trimeric holoenzyme (white, PDB: 2IAE).

Both KBTBD4-bound PP2A-A molecules adopt the same open conformation, resembling that of monomeric free PP2A-A, with an additional rotation between HEAT repeats 12-13 leading to a slightly more pronounced open arc compared to the apo structure (**Fig. 3H**) (Groves et al. 1999). This conformation differs from the closed horseshoe conformation of PP2A-A within the heterotrimeric phosphatase (**Fig. 3H**). Release of PP2A-A from the phosphatase complex allows HEAT repeats 13-15 to move outward by ∼30 degrees, which extends the KBTBD4 binding surface in PP2A-A and allows it to gain access to both Kelch domains in the KBTBD4 dimer (**Fig. 3H**). In addition to their effects on PP2A-A conformation, binding the adaptor and catalytic subunits of PP2A, PP2A-B and PP2A-C, obstructs each interaction surface of PP2A used to engage KBTBD4 (**Fig. S2F**). These findings explain why KBTBD4 can only engage free PP2A-A molecules, strengthening the notion that CRL3^KBTBD4^ acts as an orphan quality control enzyme for the major structural subunit of the PP2A phosphatase.

### Medulloblastoma mutations in *KBTBD4* disrupt protein complex quality control

We noted that MB mutations in *KBTBD4* map to the surface loop 2b-c in the Kelch domain that engages PP2A-A. These findings suggested that the very same cancer mutations that provide gain-of-function towards CoREST degradation may also impact recognition of an endogenous substrate that is a tumor suppressor. To test whether MB mutations affect recognition of PP2A-A, we immunoprecipitated KBTBD4 variants and tested for co-purifying PP2A-A. Importantly, we noted that all tested KBTBD4 MB mutants (IPR310delinsTTYML [TT], R313delinsPRR [PR], P311delinsPP [PP], G308delinsGGG [GG], I310F [IF], S309F [SF]; **Fig. S4A**) lost their ability to bind PP2A-A (**Fig. 4A**). As seen for one MB mutant, KBTBD4^PR^, loss of PP2A-A recognition disrupted ubiquitylation of PP2A-A *in vitro* and in cells, and it abolished the ability of CRL3^KBTBD4^ to instigate PP2A-A degradation (**Fig. 4B-E**). Thus, while *KBTBD4* mutations cause aberrant targeting of CoREST, they simultaneously disrupt the recognition of free PP2A-A and thus impede orphan quality control surveillance of the tumor suppressor PP2A.

**Figure 4:**
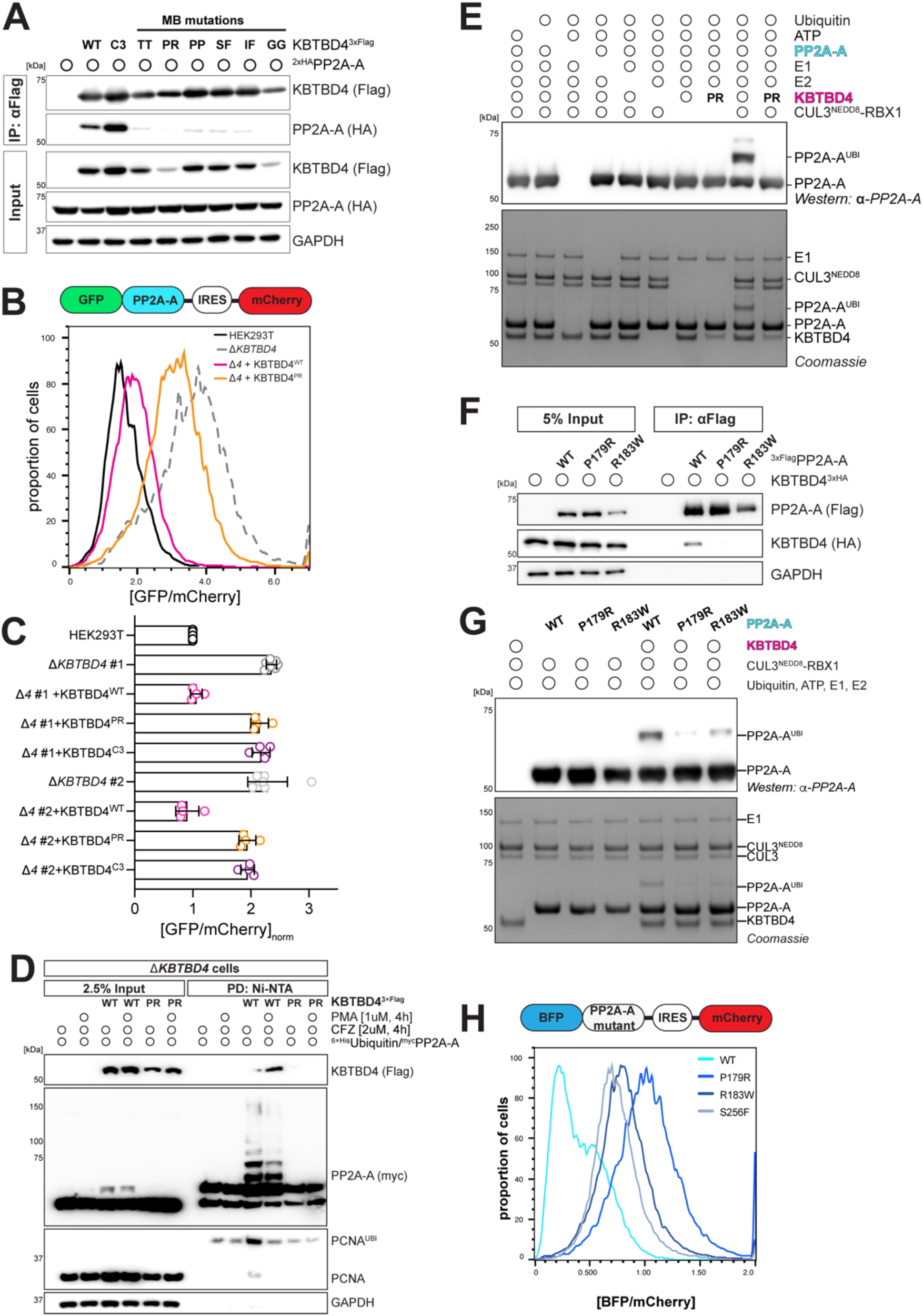
KBTBD4 MB mutations and PP2A-A cancer mutations disrupt the ligase-substrate complex formation. **A.** Flag purification of transiently expressed wild-type KBTBD4^3×Flag^ and medulloblastoma hotspot mutants to probe for binding to ^2×HA^PP2A-A. **B.** PP2A-A stability was monitored by flow cytometry through transient expression of the dual fluorescence reporter in HEK293T cells and Δ*KBTBD4* #1 with co-expression of KBTBD4^WT^ or the MB mutant KBTBD4^PR^ construct. **C.** Quantification of the stability of PP2A-A as monitored by flow cytometry after expression of the dual fluorescence reporter and indicated KBTBD4 constructs (WT, MB mutant PR, CUL3-binding mutant C3). Data was measured at least in triplicates, and the median fluorescence signal ratio of GFP over mCherry was normalized to HEK293T control. **D.** Ubiquitin conjugates were purified under denaturing conditions from Δ*KBTBD4* #1 cells, virally transduced to rescue with indicated KBTBD4 constructs, and transiently transfected to express His-tagged ubiquitin and ^myc^PP2A-A. **E.** Purified PP2A-A was ubiquitylated *in vitro* with purified KBTBD4 (WT and MB cancer mutant PR), CUL3^NEDD8^-RBX1, UBE1, UBE2D3 and fluorescently labeled ubiquitin. Representative result of n=2 independent experiments. **F.** Flag purification of transiently expressed ^3×Flag^PP2A-A (wild-type and cancer mutants P179R and R183W) to probe for binding to KBTBD4^3×HA^. **G.** Purified PP2A-A (WT, and cancer mutants P179R and R183W) was ubiquitylated *in vitro* with purified KBTBD4, CUL3^NEDD8^-RBX1, UBE1, UBE2D3 and fluorescently labeled ubiquitin. Representative result of n=2 independent experiments. **H.** Quantification of the stability of wild-type PP2A-A and indicated mutants as monitored by flow cytometry after viral transduction of a dual fluorescence reporter. Data representative of n=3 independent experiments.

Similar to *KBTBD4*, *PPP2R1A*, the gene encoding PP2A-A, is mutated in cancer (Taylor et al. 2019; O’Connor et al. 2020). The commonly mutated residues in PP2A-A, P179 and R183, are within HEAT repeat 5 and point directly towards KBTBD4 (**Fig. 3F**). Accordingly, the uterine-cancer variants PP2A-A^P179R^ and PP2A-A^R183W^ lost their ability to bind KBTBD4 (**Fig. 4F**). Also *in vitro,* mutation of P179 and R183 in PP2A-A impaired their ubiquitylation by CRL3^KBTBD4^ (**Fig. 4G**). In line with these observations, the P179R and R183W variants of PP2A-A were stabilized in cells to a similar extent as WT PP2A-A in cells lacking *KBTBD4* (**Fig. 4H**). Thus, PP2A-A mutants in cancer strongly impair protein complex quality control through CRL3^KBTBD4^.

### Endogenous phospho-regulated pathways are affected by loss of KBTBD4

PP2A is a major Ser- and Thr-directed phosphatase that modulates pathways important for cellular fitness and is often dysregulated in cancer (Brautigan and Shenolikar 2018). To assess whether elevated levels of PP2A-A impact cellular fitness, we designed a competition assay to compare cells that overexpressed wild-type or KBTBD4-binding deficient PP2A-A variants to cells that expressed only endogenous PP2A-A levels (**Fig. 5A**). We observed a slight reduction in cell viability when we overexpressed wild-type PP2A-A, indicating that increased levels of PP2A-A reduce cellular fitness. Importantly, the P179R and R183W variants of PP2A-A imparted a very strong fitness defect on cells. The inability of these cancer variants to form trimeric PP2A holoenzymes (Taylor et al. 2019; O’Connor et al. 2020) together with their failure to undergo CRL3^KBTBD4^-dependent orphan quality control, seem to increase the cellular pool of unassembled PP2A-A in a manner that negatively impacts cell viability.

**Figure 5:**
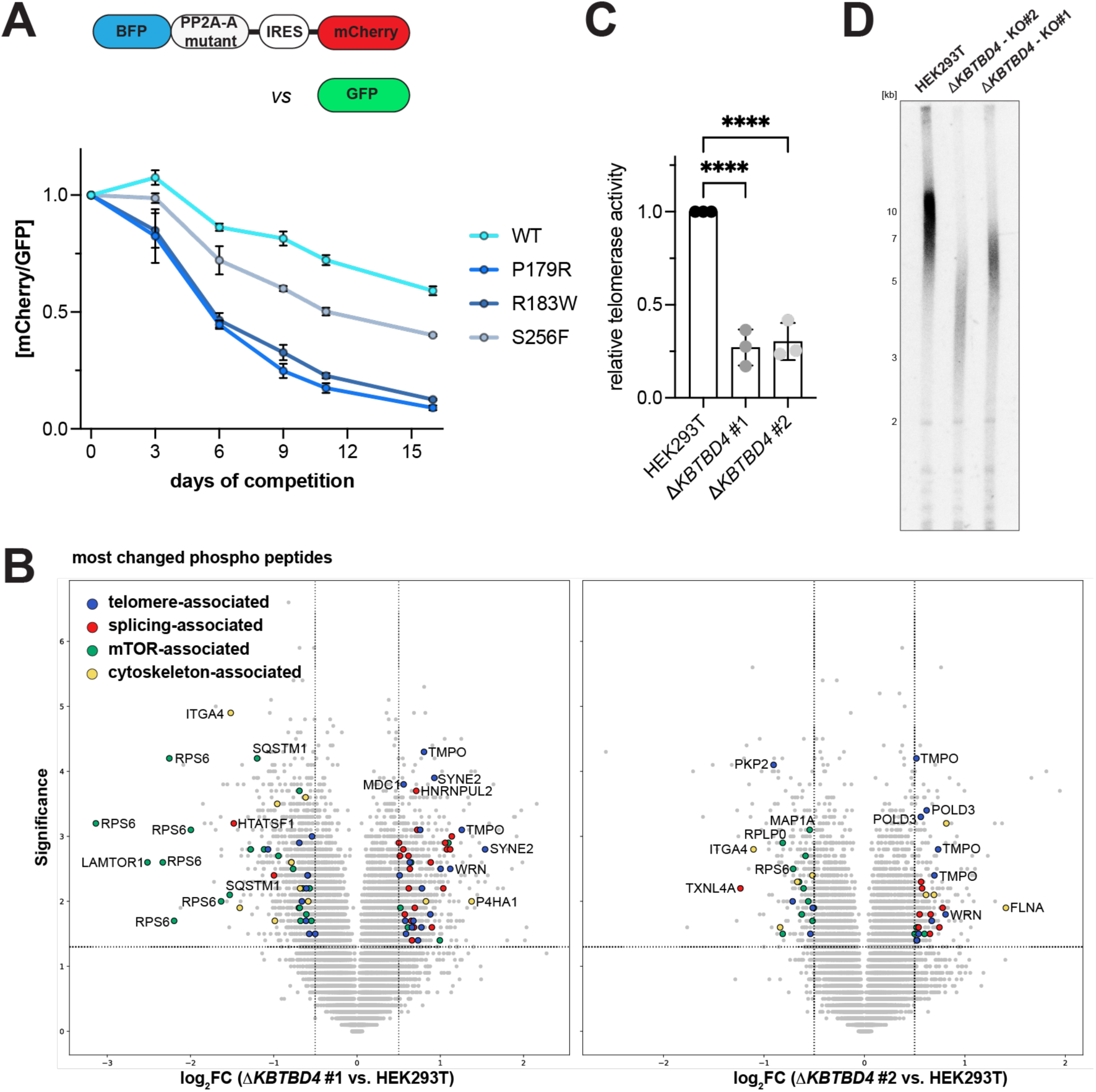
Elevated cellular PP2A-A levels disrupt phospho signaling and correlate with impaired telomerase activity and reduced telomere length. **A.** Relative cellular fitness of cells overexpressing PP2A-A (WT or indicated mutants) was determined by flow cytometry following viral transduction with PP2A-A dual fluorescence reporter constructs or a GFP control construct. Fitness was inferred from changes in the mCherry/GFP ratio during long-term passaging under selection, normalized to day 0. **B.** Phospho-proteomic analysis of Δ*KBTBD4* cell lines #1 and #2 compared to wild-type HEK293T cells. Volcano plots display the log2 fold change of phospho-peptide abundance versus significance (-log_10_(p-value)); phospho peptides meeting the indicated thresholds (*p* ≤ 0.05; log2FC ≥ ±0.5) and are significantly altered in both cell lines are highlighted and grouped by pathway. **C.** Telomerase activity in HEK293T and Δ*KBTBD4* cell lines was quantified by RQ-TRAP, a quantitative PCR-based modification of the TRAP assay, and is shown as relative activity normalized to HEK293T cells; data represent biological triplicates. **D.** Length of telomeres was assessed by telomere restriction fragment analysis of HinfI/RsaI-digested genomic DNA from HEK293T and Δ*KBTBD4* cell lines. Telomeric DNA was detected using a radiolabeled telomeric probe; data shown are n=1.

Does loss of KBTBD4-function and the ensuing rise of free PP2A-A impact cellular phospho signaling? To address this, we used mass spectrometry to measure changes in the phospho-proteome of Δ*KBTBD4* cells compared to their wild-type counterparts (**Fig. S4C-E**). We found that *KBTBD4* deletion changed phosphorylation patterns of mTOR-, spliceosome-, and telomerase-associated proteins (**Fig. 5B, Fig. S4D**) Intriguingly, telomere maintenance has been identified as an enhanced molecular signature within the KBTBD4-driven MB subgroup 3 (Cavalli et al. 2017), and telomerase activity is regulated by PP2A (Li et al. 1997). We therefore analyzed the effects of loss of KBTBD4 on telomerase activity in a quantitative PCR-based telomeric repeat amplification (qTRAP) assay (Wege et al. 2003) and in Southern blots measuring telomere length. Telomerase activity in Δ*KBTBD4* cells was significantly reduced (**Fig. 5C**), and Southern blots showed a coherent telomere length reduction compared to wild-type cells (**Fig. 5D**). Consistently, DepMap data reveals a deleterious effect of mutations in the catalytic subunit of telomerase, *TERT*, in *PPP2R1A* deletion (**Fig. S4F**, 25Q3 release). Together, these findings showed that inactivation of KBTBD4, as observed in medulloblastoma, abolishes orphan quality control of PP2A, thereby disrupting PP2A- and telomere-regulation that were long known to be dysregulated in cancer.

## Discussion

Hotspot mutations in the E3 ligase adaptor *KBTBD4* have been shown to act as gain-of-function events in medulloblastoma, while their effect on the endogenous function of the ligase remained unclear. Here, we show that these mutations in *KBTBD4* also trigger a loss-of-function phenotype impairing the role of KBTBD4 in orphan quality control of the scaffolding subunit of the phosphatase PP2A. Mutations in KBTBD4 hence drive cancer both by impairing tumor suppressor function through loss of PP2A maintenance, as well as a simultaneous gain-of-function oncogenic event degrading HDAC-associated complexes. The precise assembly of protein complexes and their balanced stoichiometry is essential for cell fate decisions (Padovani et al. 2022). Perturbations in the stoichiometric balance of protein complex components impair the function of important cellular regulators, resulting in misregulation of downstream developmental pathways and the emergence of developmental disorders and cancer. Orphan quality control eliminates overabundant subunits, while protein complex quality control corrects defective protein assemblies. Both are dependent on the ubiquitin-proteasome system, and both are dysregulated in cancer (Mark et al. 2023). The E3 ligase CRL3^KBTBD4^, found mutated in medulloblastoma, plays a critical role in orphan quality control of PP2A-A, the core architectural subunit of the PP2A phosphatase. Thus, rather than solely driving tumorigenesis through degrading CoREST subunits, mutations in *KBTBD4* have complex consequences that combine gain-of-function CoREST degradation with loss-of-function phosphatase regulation.

PP2A is one of the most abundant enzymes, making up ∼1% of total cellular protein in certain tissues, and being responsible for ∼40% of all dephosphorylation events in humans (Fowle et al. 2019). CLR3^KBTBD4^ ubiquitylates free PP2A-A that is not incorporated into PP2A complexes (**Fig. 2B**). The cryo-EM structure shows that KBTBD4 selectively recognizes an extended conformation of PP2A-A that is only observed upon dissociation of the catalytic subunit, PP2A-C (**Fig. 3A, H**), and how residues commonly mutated in *KBTBD4* in medulloblastoma mediate recognition of free PP2A-A (**Fig. 3C, D**). While for CoREST degradation, KBTBD4 indel mutations generally outperformed single point substitutions (Xie et al. 2025), we already observe almost complete loss of binding for the S309F and I310F KBTBD4 variants (**Fig. 4A**). Despite the overlap of the KBTBD4 surface for UM171/CoREST (Yeo et al. 2025) and PP2A-A binding, the detailed molecular mechanisms differ and endogenous PP2A-A degradation is not obviously connected to gain-of-function degradation of CoREST components. Strikingly, residues in PP2A-A that contact KBTBD4 are also commonly mutated in uterine cancer (**Fig. 3F**). Our functional and structural data together provide the molecular basis for how tumor mutations additionally inhibit orphan quality control of PP2A-A through CRL3^KBTBD4^, highlighting an important role of this quality control system in preventing oncogenesis.

PP2A-A is produced in excess over PP2A-C to efficiently form the PP2A-AC core, which in turn must be bound by inhibitors such as SET (Li et al. 1996) to prevent rouge phosphatase activity. When PP2A is disassembled, the PP2A inhibitor CIP2A associates with PP2A-BC to release free PP2A-A (Pavic et al. 2023). Orphan quality control through CRL3^KBTBD4^ supports the process through the subsequent removal of free PP2A-A. Accordingly, we observe accumulation of monomeric PP2A-A in Δ*KBTBD4* cells (**Fig. 2C**). Analogous to mutations in KBTBD4, alterations in the heterotrimeric PP2A phosphatase are also implicated in oncogenesis. The most prevalent PP2A-A mutations, P179R and R183W, disrupt both PP2A holoenzyme assembly (Taylor et al. 2019; O’Connor et al. 2020) as well as PP2A-A orphan quality control mechanisms. The impaired quality control could result from weakened binding of mutant PP2A-A to KBTBD4, either due to direct disruption of the binding interface or mutation-induced conformational changes in PP2A-A (Taylor et al. 2019), resulting in markedly elevated levels of free PP2A-A (**Fig. 4H**). Notably, PP2A-A cancer mutations appear to preferentially target the shared KBTBD4/PP2A-B interface. Our work proposes that the oncogenic effects of these mutations arise not only from impaired PP2A holoenzyme formation and reduced enzymatic activity, but also from dysregulated PP2A-A orphan quality control.

Phospho-proteomics showed that loss of PP2A-A orphan quality control results in changes of phosphorylation, indicative of altered PP2A activity (**Fig. 5B**). This was particularly evident in two pathways with links to MB: mTOR signaling, associated with the SHH subgroup (Wu et al. 2017); and telomere maintenance, identified as a vulnerability in MB subgroup 3 (Cavalli et al. 2017). Phosphorylation of telomere-associated or regulatory proteins such as the helicase WRN and the crossover junction specific endonuclease MUS81 increased upon CRL3^KBTBD4^ loss, and we find telomerase activity impaired, and consistently telomere length reduced in these cells. As WRN helicase inhibitors are moving towards clinical trials (Rodríguez Pérez et al. 2024), our findings could therefore help stratify patients that could benefit from such molecules.

Overall, our findings suggest that KBTBD4 mutations in medulloblastoma impair PP2A regulation and function. This loss-of-function phenotype accompanies the gain-of-function CoREST degradation to drive tumorigenesis, showing that a single point mutation in a critical E3 ligase can lead to substantial rewiring of critical signaling networks in cancer.

---

**Figure S1:**
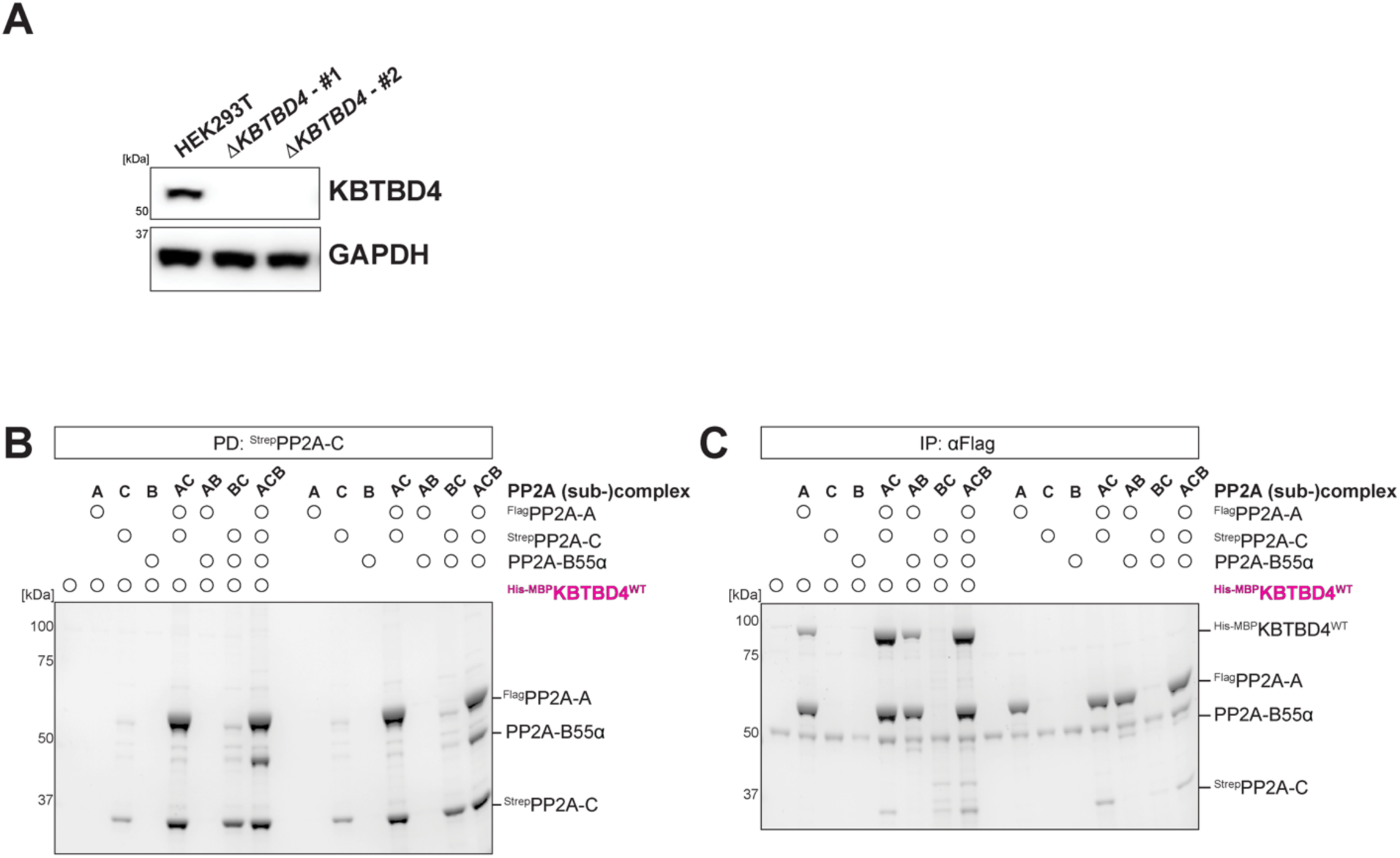
Generation of KBTBD4 knockout cell lines and analysis of KBTBD4 binding to PP2A subcomplexes. **A.** The clonal knockout cell lines Δ*KBTBD4* #1and #2 were generated from parental HEK293T cells and KBTBD4 protein levels were analyzed by western blot. **B** and **C.** Monomeric PP2A subunits PP2A-^3×Flag^Aα/^Strep^Cα/B55α were expressed as individual proteins, in PP2A subcomplexes PP2A-AC/AB/CB, and the heterotrimeric PP2A-ACB holoenzyme in insect cells. PP2A extracts were mixed with ^6×His-MBP^KBTBD4 extract and affinity purified as indicated to test for PP2A complex formation in extracts and binding of PP2A-C and PP2A-A to KBTBD4.

**Figure S2:**
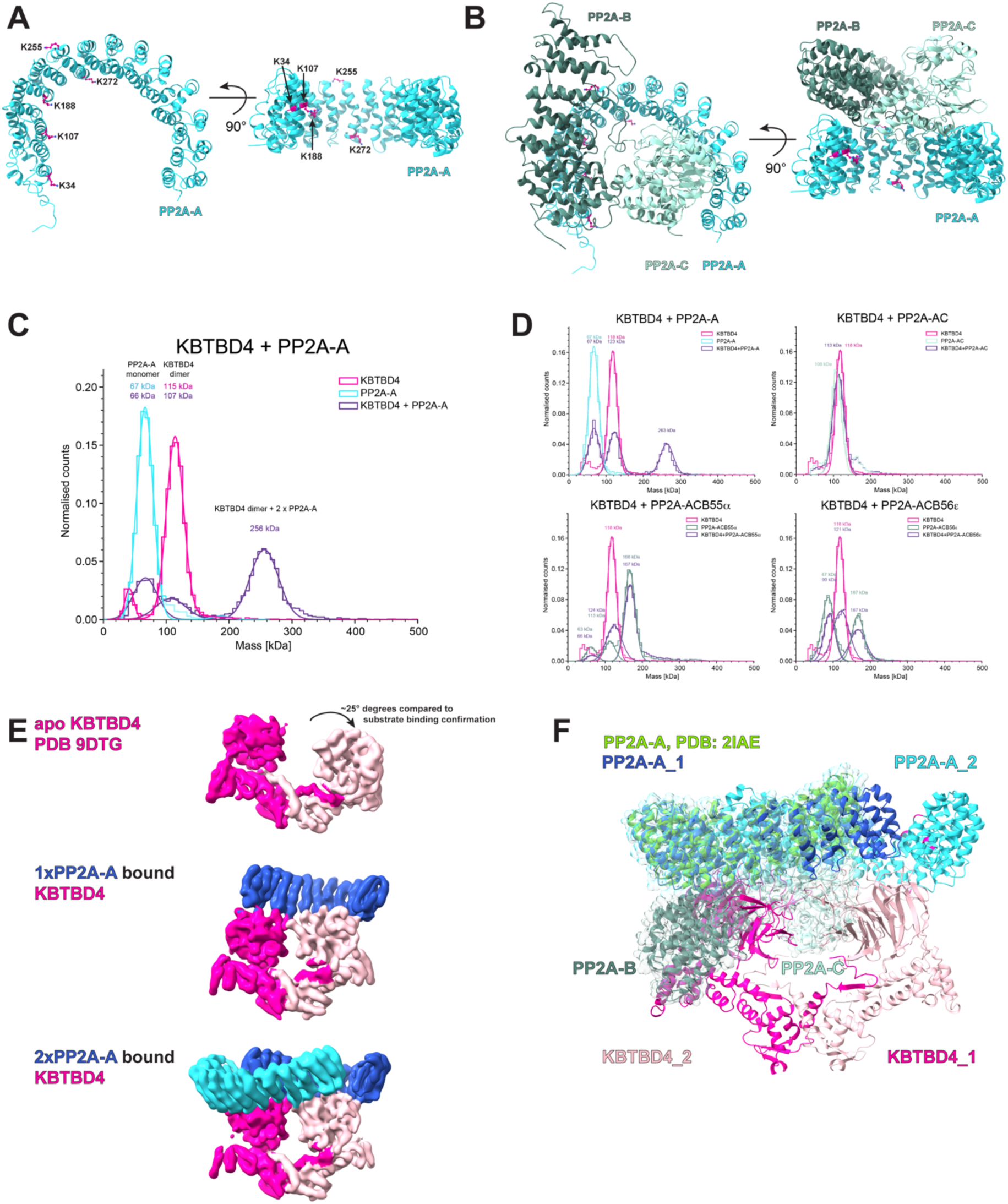
Structural and biophysical analysis of KBTBD4 interaction with PP2A-A. **A** and **B.** Ubiquitylation sites in PP2A-A were determined by mass spectrometry and mapped onto PP2A-A (PDB: 2IAE). **C** and **D.** Mass photometry measurements of KBTBD4 and PP2A-A or PP2A (sub-) complexes alone or mixed at a 1:1 molar ratio. Representative mass distributions are shown, with molecular masses determined using a Refeyn OneMP instrument calibrated with a BSA standard. **E.** Low pass filtered cryo-EM maps of apo KBTBD4 (PDB: 9DTG), a 2:1 complex of KBTBD4:PP2A-A (EMD-56547), and a 2:2 complex of KBTBD4:PP2A-A (EMD-56545). **F.** Overlay of the atomic models of KBTBD4 or PP2A-CB bound PP2A-A (PDBs 2IAE vs 28JO).

**Figure S3:**
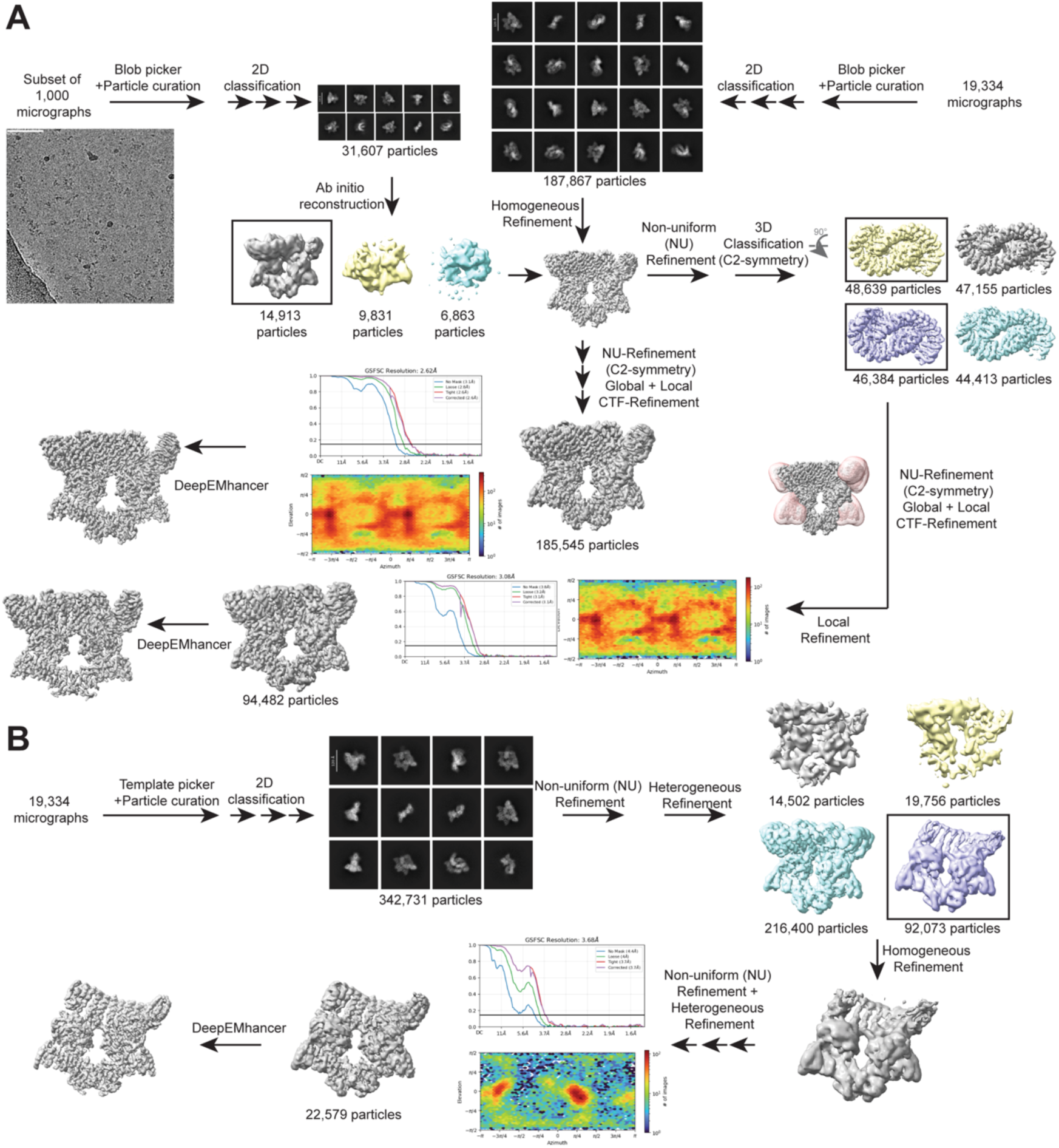
Cryo-EM processing scheme of PP2A-A bound to KBTBD4. **A.** Cryo-EM processing scheme of two copies of PP2A-A bound to dimeric KBTBD4. The micrograph is representing the collected 19,334 micrographs of this dataset of which a subset of 1,000 micrographs was used to generate the *ab initio* model subsequently used for the full dataset. Scalebars on 2D classes correspond to 120 Å. Data processing was performed in CryoSparc v4.7.1. A final 3D reconstruction was obtained at 2.62 Å by the gold-standard Fourier shell correlation of 0.143 with 185,545 particles. Additionally, local refinement was performed with a mask indicated in transparent pink covering regions of lower resolutions to aid model building and resulted in a final resolution of 3.08 Å with 94,482 particles. **B.** Cryo-EM processing scheme of one copy of PP2A-A bound to dimeric KBTBD4. To obtain a map corresponding to dimeric KBTBD4 binding to a single PP2A-A moiety, the full dataset of A was reprocessed in CryoSparc v4.7.1 and selecting for 2:1 complex during heterogeneous refinement. Using 22,579 particles, a final 3D reconstruction of 3.68 Å was obtained.

**Figure S4:**
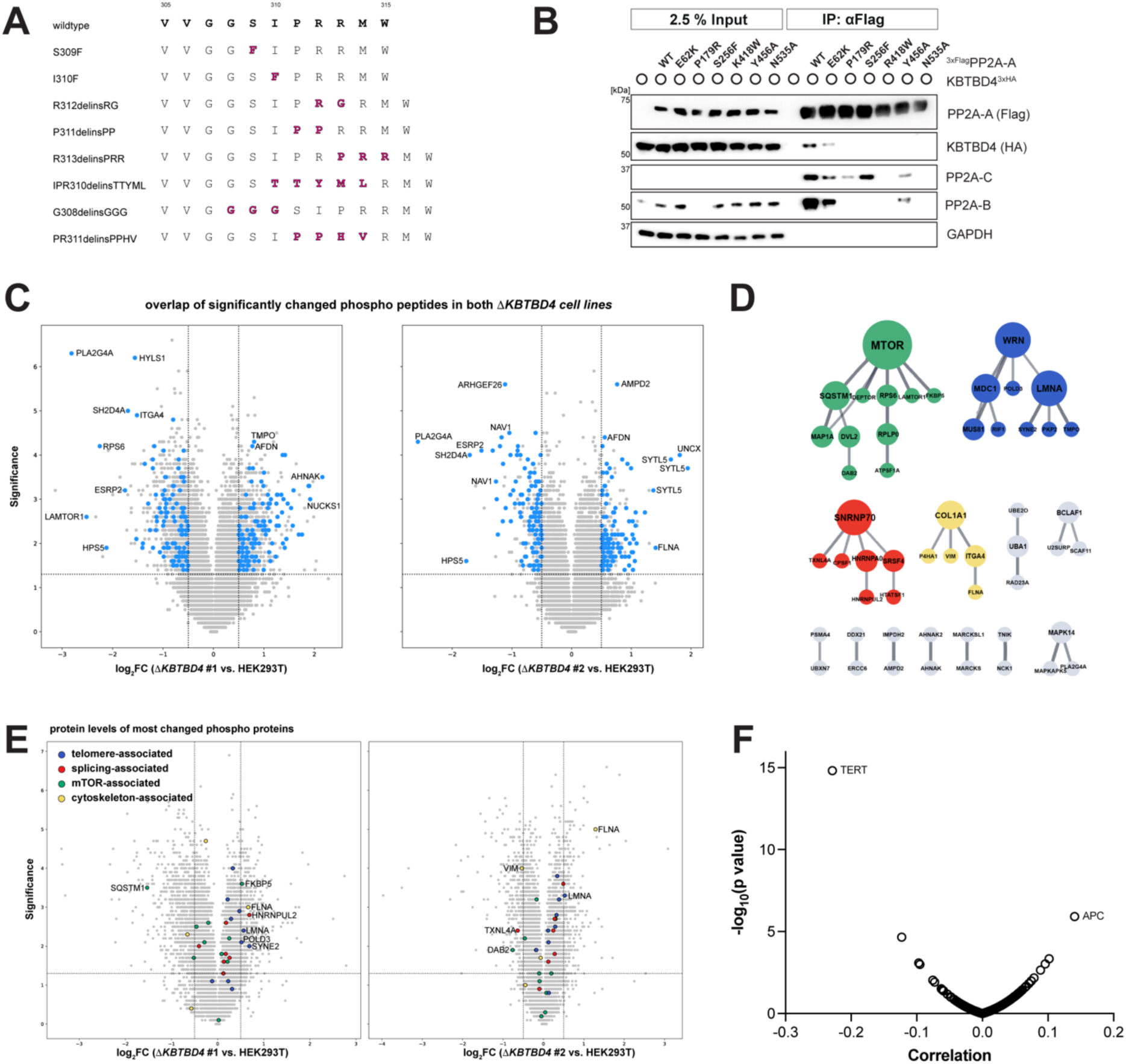
Functional consequences of KBTBD4 mutation and deletion on PP2A-A binding and phospho signaling. **A.** Amino acid sequences of medulloblastoma hotspot mutations in KBTBD4. **B.** Flag purification of transiently expressed ^3×Flag^PP2A-A (wildtype and indicated) to probe for binding to KBTBD4^3×HA^. **C.** Same volcano plots as Fig. 5B, highlighting in blue the phospho peptides that were meeting the significance cutoff (log_2_FC > 0.5 or < -0.5; significance ≥ 1.3) in both Δ*KBTBD4* cell lines, and were used for clustering analysis. **D.** String-DB and Cytoscape analysis of the changed phospho peptides as highlighted in C to identify most changed pathways (Shannon et al. 2003). **E.** Same volcano plots as Fig. 1B, highlighting the relative abundance of the proteins with the most changed phospho peptides. **F.** Plot of DepMap correlation data for the effect of hotspot mutations in *PPP2R1A* deletion (release 25Q3).

## Materials and Methods

### Mammalian Cell Culture

Human embryonic kidney (HEK) 293T cells were maintained in DMEM + GlutaMAX (Gibco, 10566-016) with 10% fetal bovine serum ([FBS]; VWR, 89510-186). Cells were purchased from the UC Berkeley Cell Culture Facility and ATCC, and routinely tested for mycoplasma contamination using a Mycoplasma PCR Detection kit (abmGood, G238). All cell lines tested negative for mycoplasma. Plasmid transfections were performed using polyethylenimine (PEI or PEI MAX; Polysciences 23966-1 and 24765) at a 1:6 ratio of DNA (in μg) to PEI (in μL at a 1 mg mL^-1^ stock concentration) or Lipofectamine 3000 transfection reagent (Thermo Fisher, L3000008) per the manufacturers’ instructions. Lentiviruses were produced in HEK293T cells by co-transfection of lentiviral and packaging plasmids using Lipofectamine 3000. Virus-containing supernatants were collected after 48 h, spun down, aliquoted and stored at -80°C for later use. For lentiviral transductions, 2×10^5^ cells were seeded into 6-well plates and transduced by addition of lentiviral particles and 6 μg mL^-1^ polybrene (Sigma-Aldrich, TR-1003). HEK293T transduced cells were drug-selected 24 h after infection with puromycin (at 1 μg mL^-1^; Sigma-Aldrich, P8833), or blasticidin (at 7.5 μg mL^-1^; Thermo Fisher, A1113903).

### Plasmids

The list of all constructs used in this study are provided in Supplementary Table XX. Most cloning was performed by restriction site cloning or Gibson assembly using HiFi DNA Assembly master mix (NEB, E2621L).

### Antibodies

The following commercially available antibodies were used in this study: anti-KBTBD4 (Novus Biologicals, NBP1-88587), Anti-FLAG (Sigma-Aldrich, Clone M2, F1804), anti-DYKDDDDK-Tag (Cell Signaling Technology [CST], 2368), anti-HA-Tag (CST, 3724), anti-CUL3 (Bethyl Laboratories, A301-110A), anti-PP2A-A (CST, 2041), anti-PP2A-C (CST, 2038), anti-PP2A-B (CST, 2290), anti-GAPDH (CST, 2118), anti-Myc-Tag (CST, 2278), anti-PCNA (CST, 2586).

### Generation of CRISPR-Cas9 Genome Edited **Δ***KBTBD4* Cell Lines

The Δ*KBTBD4* cell lines of this study were generated from HEK293T cells. Guide RNAs (sgRNAs) were designed using the IDT online resource to target the genomic locus of the KBTBD4 gene on chromosome 11 within introns 1 (5’-ACACCATCTTGAGTAACCTG-3’) and 3 (5’-TCAGGACAAGCTTGAAGGTA-3’), removing exons 2 and 3. DNA oligonucleotides for the designed sgRNAs and their complementary sequences were phosphorylated (New England Biolabs [NEB], M0201), annealed, and ligated (NEB, M0202) into pX330 (Cong et al. 2013). Low-passage HEK293T cells were cultured in a 12-well plate and transfected at 50% confluency with 1 μg of each pX330 plasmid (targeting intron 1 and 3, respectively) and Mirus TransIT-293T reagent (Mirus Bio, MIR2705). After 48 h, cells were expanded to a 6-well format and analyzed for successful bulk editing by PCR. After another 48 h, individual clones were seeded in 96-well plates for single-cell selection. After two weeks, individual homozygous clones were expanded, screened by PCR and DNA sequencing and confirmed by western blotting.

### Immunoprecipitation and Mass Spectrometry (IP-MS)

Anti-Flag immunoprecipitations (IPs) for mass spectrometry analysis were performed from extracts of 20×15 cm plates of HEK293T cells transiently expressing KBTBD4^3×Flag^. Drug treatments were performed with MLN4924 (Cayman Chemicals, 15217) and Carfilzomib (CFZ; Sellek Chemicals, S2853) at 5 μM and DMSO as vehicle control for 8 h before harvesting. Cells were lysed in lysis buffer (20 mM HEPES pH 7.4, 100 mM NaCl, 10 mM NaF, 1 mM Na_3_VO_4_, 0.2% NP40, 1× cOmplete protease inhibitor cocktail [Roche 11836170001], benzonase [Sigma-Aldrich, 70746-4] and extracts were clarified by centrifugation at 21,000×g and bound to ANTI-FLAG M2 affinity resin (Sigma-Aldrich, A2220) for 2 h at 4°C. Immunoprecipitates were then washed 4× and eluted 3× at 30°C with 0.5 mg mL^-1^ of 3×Flag peptide (Sigma, F4799) buffered in 1× PBS plus 0.1% Triton X-100. Elutions were pooled and precipitated overnight at 4°C with 20% trichloroacetic acid. Spun down pellets were washed 3× with an ice-cold acetone and 0.1 N HCl solution, dried, resolubilized in 8 M urea buffered in 100 mM Tris pH 8.5, reduced with TCEP (Sigma-Aldrich, C4706) at a final concentration of 5 mM for 20 min, alkylated with iodoacetamide (Thermo Fisher, A39271) at a final concentration of 10 mM for 15 min, diluted 4-fold with 100 mM Tris pH 8.5, and digested with 0.5 mg mL^-1^ of trypsin (Promega, v5111) supplemented with CaCl_2_ at a final concentration of 1 mM overnight at 37°C. Trypsin-digested samples were submitted to the Vincent J. Coates Proteomics/Mass Spectrometry Laboratory at UC Berkeley for analysis. Peptides were processed using multidimensional protein identification technology (MudPIT) and ran on a LTQ XL linear ion trap mass spectrometer. Protein spectral counts were normalized to bait counts, multiplied by 100, added 1 ((spectral counts_protein_/spectral counts_KBTBD4_) × 100 +1). Changes induced by drug treatment were quantified as log_2_ fold changes of MLN4924-or CFZ-treated samples relative to DMSO control and visualized by plotting log_2_(MLN4924/DMSO) against log_2_(CFZ/DMSO).

### Proteomics and Phospho-Proteomics

For proteomics and phospho-proteomics analysis wild-type HEK293Ts, Δ*KBTBD4* knockout cell lines #1 and #2 were cultured in DMEM (Life Technologies) supplemented with 10% (v/v) fetal bovine serum (FBS, Thermo Fisher Scientific), 2 mM L-glutamine (Lonza), 100 U/mL penicillin (Lonza) and 0.1 mg mL^-1^ streptomycin (Lonza). When cells were confluent, culturing media was refreshed for 48 h in n=3 biological replicates. Cells were washed twice with PBS (Gibco), collected in LoBind tubes (Eppendorf) in 5% SDS buffer supplemented with Protease (Roche) and Phosphatase (Roche) inhibitors. Cells were lysed by sonication and lysates were cleared by centrifugation and protein concentrations were determined using Pierce BCA assay (Thermo Scientific). 300 μg of protein from each condition were aliquoted into LoBind tubes and the STRAP (Protifi) protocol was followed according to manufacturer’s instructions with slight modifications. The protein was reduced with 5 mM DTT at 55°C for 15 min, alkylated (final concentration 20 mM IAA or CAA) at room temperature for 10 min and then acidified (12% phosphoric acid). Protein was then trapped into S-trap columns and washed with S-trap binding/wash buffer (100 mM TEAB in 90% methanol) by centrifugation at 4,000×g for 30 s. Cleaned proteins then went through tryptic digestion (Trypsin (Thermo Scientific) in 50 mM TEAB) overnight at 37°C or 2 h at 47°C. Digested peptides were sequentially eluted in 80 μL of 50 mM TEAB (Sigma-Aldrich), 0.2% formic acid (FA, Sigma-Aldrich) and 50% acetonitrile (ACN, Sigma-Aldrich). Elutions were pooled and dried down. Dried and cleaned peptides were reconstituted in 110 μL of 100 mM TEAB. TMT-16-plex reagent (Thermo Scientific) was brought to room temperature and reconstituted in 20 μL of anhydrous acetonitrile. TMT reagent solution was transferred to the peptides and incubated on a thermoshaker (Eppendorf) at 400 rpm at room temperature for 1 h. The reaction was quenched by the addition of 8 μL of 5% Hydroxylamine for 30 min at room temperature. TMT-labelled peptides were pooled together and dried down in the speed vac for downstream processing.

Pooled peptides were separated by basic reverse phase chromatography fractionation on a C_18_ column with flow rate at 200 μL/min with two buffers: buffer A (10 mM ammonium formate, pH 10) and buffer B (80% ACN, 10 mM ammonium formate, pH 10). Peptides were resuspended in 100 μL of buffer A (10 mM ammonium formate, pH 10) and resolved on a C_18_ reverse phase column by applying a non-linear gradient of 7-40%. A total of 80 fractions were collected and concentrated into 24 fractions for the proteomics analysis and 12 for phospho-proteomics enrichment. For proteomics, only 5% samples were used and remaining 95% were used for the phospho-proteomics analysis. For phospho-peptide enrichment Ni-NTA (nitrilotriacetic acid) superflow agarose beads were used. The nickel from the beads were stripped with 100 mM EDTA and incubated in an aqueous solution of 10 mM iron-(III)-chloride (FeCl_3_). Dried peptide fractions were reconstituted to a concentration of 0.5 μg/μL in 80% ACN/0.1% TFA. Peptide mixtures were enriched for phosphorylated peptides with 10 μL IMAC beads for 30 min with end-to-end rotation. Enriched IMAC beads were loaded on Empore C_18_ silica packed stage tips. Stage tips were equilibrated with methanol followed by 50% ACN/0.1% FA then 1% FA. The beads with enriched peptides were loaded onto C_18_ stage tips and washed with 80% ACN/0.1% TFA. Phosphorylated peptides were eluted from IMAC beads with 500mM dibasic sodium phosphate, pH 7.0. These peptides were washed with 1% FA before elution using 50% acetonitrile in 0.1% FA. The peptides were then dried by SpeedVac and stored at -20°C until mass spectrometry analysis. Enriched phospho-peptides and peptides were analyzed on an Orbitrap Ascend Tribrid mass spectrometer interfaced with Dionex Ultimate 3000 nanoflow liquid chromatography system.

For proteomics analysis peptides were separated on an analytical column (Acclaim PepMap RSLC C18, 75 μm × 50 cm, 2 μm, 100 Å) at a flow rate of 300 nL/min, using a step gradient of 2-7% solvent B (90% ACN/0.1% FA) for the first 6 min, followed by 7-18% up to 89 min, 18-27% up to 89-114 min and 27-35% to 114-134 min. The total run time was set to 155 min. The mass spectrometer was operated in a data-dependent acquisition mode in SPS MS3 (FT-IT-HCD-FT-HCD) method. A survey full scan MS (from m/z 350-1500) was acquired in the Orbitrap at a resolution of 120,000 at 200 m/z. The AGC target for MS1 was set as 4×105 and the ion filling time as 50 ms. The precursor ions for MS2 were isolated using a Quadrupole mass filter at a 0.7 Da isolation width, fragmented using a normalized 30% HCD of ion routing multipole and analyzed using ion trap. The top 10 MS2 fragment ions in a subsequent scan were isolated and fragmented using HCD at a 55% normalized collision energy and analyzed using an Orbitrap mass analyzer at a 60,000 resolution, in the scan range of 100-500 m/z.

For phospho-proteomics, LC setting is same. The mass spectrometer was operated in a data-dependent acquisition mode in MS2 (FT-HCD) method. A survey full scan MS (from m/z 350-1500) was acquired in the Orbitrap at a resolution of 120,000 at 200 m/z. The AGC target for MS1 was set as 4×105 and the ion filling time as 251 ms. The monoisotopic precursor selection was used. The precursor ions for MS2 were isolated at 1.6 Da isolation width, fragmented using a normalized 30% HCD. The AGC targets for MS2 was used as 50,000 and ion injection time were set as 123 ms. Orbitrap resolution for MS2 were set as 60,000.

The proteomics raw data were searched using SEQUEST HT search engines with Proteome Discoverer 3.0 (Thermo Fisher Scientific). The parameter used for the proteomics analysis: MS1 tolerance 10 ppm, MS2 mass tolerance 0.6, enzyme: trypsin, Mis-cleavage: -2, fixed modification: carbamidomethylation of cysteine residues and TMT of lysine and N-terminal, dynamic modification: oxidation of methionine. The data were filtered for 1% protein level FDR. The phospho-proteomics raw data were searched using SEQUEST HT search engines with Proteome Discoverer 3.0 (Thermo Fisher Scientific). The following parameters were used for searches: Precursor mass tolerance 10 ppm, fragment mass tolerance 0.02, enzyme: trypsin, Mis-cleavage: -2, fixed modification: carbamidomethylation of cysteine residues and TMT of lysine and N-terminal, dynamic modification: oxidation of methionine and phosphorylation of serine, threonine and tyrosine. The data were filtered for 1% PSM and peptide level. A two-tailed *t*-test with 250 randomizations was performed in Perseus software (v1.6.15.0) to assess statistical significance between groups. Site localization probabilities for phosphorylation events were calculated using the ptmRS node. Analysis of protein abundance was plotted in Fig. 1B (for PP2A subunits) and Fig. S4E (for total protein abundance of proteins with most changed phospho peptides). Analysis of phospho peptides was plotted in Fig. 5B (highlighting of most changed phospho peptides by clustering analysis) and Fig. S4C (highlighting of phospho peptides that are significantly changed in both Δ*KBTBD4* cell lines, as used for clustering analysis).

### Global Protein Stability Flow Reporter Assay

To assess protein stability of the PP2A subunits, PP2A-A/C/B sequences were cloned as GFP-fusions with an IRES-separated mCherry for internal normalization in a pCS2 vector. HEK293T or Δ*KBTBD4* cells were plated at 2×10^5^ cells per 6-well for transfection. After 24 h, Lipofectamine 3000 (Thermo Fisher, L3000008) transfections were performed in a 6-well format according to the manufacturer’s instructions. Briefly, 125 μL of OptiMEM (Gibco, 31985070) was added to two Eppendorf tubes A and B. 200 ng of reporter plasmid DNA and 5 μL P3000 reagent were diluted in tube A, and 5 μL Lipofectamine 3000 reagent was diluted in tube B. For the rescue experiments, 200 ng of the reporter plasmid were supplemented with 1.8 μg of either control vector pCS2+, or the KBTBD4 constructs cloned in pCS2. Tube A was added to tube B, incubated at RT for 10 min, and 70 μL of the mix were pipetted onto the cells per well. Cells expressed the constructs for 48 h before analysis. Drug treatments were performed at the indicated concentrations for 6 h before analysis and compared to DMSO treatment. For flow cytometry, cells were trypsinized, washed, and resuspended in flow buffer, to be analyzed on a BD LSR Fortessa instrument. The data was analyzed with FlowJo (BD Biosciences, v10.10.0) and for quantifications, the median fluorescence signal ratio of GFP over mCherry was normalized to controls and plotted with GraphPad Prism (v10.6.1) to assess stability of the protein.

### Small-Scale Co-Immunoprecipitations from Mammalian Cells

HEK293T were seeded at 1.2×10^6^ cells in a 10 cm plate and transfected after 24 h with a total of 3 μg plasmid DNA (typically a ratio of 2:1 of bait to prey constructs) and PEI at a ratio DNA/PEI of 1:6. After 48 h of expression, the cells were harvested with PBS, and collected by centrifugation at 300×g for 5 min. Harvested cell pellets were either flash frozen in liquid nitrogen and stored at -80°C, or immediately used for Flag-affinity purifications. For affinity purifications, pellets were lysed in 500 μL freshly prepared IP buffer (20 mM HEPES pH 7.4, 100 mM NaCl, 10 mM NaF, 1 mM Na_3_VO_4_, 2 mM phenanthroline, 0.2% NP40, protease inhibitor cocktail) + 0.5 μL benzonase (Sigma-Aldrich, 70746-4) per sample for 20 min and clarified by centrifugation at 4°C for 20 min at 21,000×g. Supernatants were normalized to volume and protein concentration using the commercially available Pierce 660 Protein Assay reagent (Thermo Fisher, 22660). Next, 5% of the supernatant was removed as input sample and diluted in 2× urea sample buffer (USB) + 5% β-mercaptoethanol (BME). The remaining sample was added to pre-equilibrated ANTI-FLAG M2 affinity slurry (Sigma-Aldrich, A2220) and rotated at 4°C for 1-2 h. For the lysate from a 10 cm plate, a volume of 30 μL affinity resin slurry was used. After incubation, the beads were washed 3× with 1 mL lysis buffer and eluted in USB. The samples were used for subsequent SDS-PAGE and Western blot analysis. The gels were run with standard amounts of 25% precipitate, compared to 2.5% or 5% relative input.

### Small-Scale Co-Immunoprecipitations from Insect Cells

Co-immunoprecipitations to assess interactions between KBTBD4 and various PP2A (sub-)complexes were performed using recombinantly expressed proteins from insect cells. In brief, Hi5 cells were infected with baculoviruses encoding His-MBP-tagged KBTBD4 and NUDCD3 (to support KBTBD4 folding) or different PP2A subcomplexes (Flag-tagged PP2A-A, Strep-tagged PP2A-C, untagged PP2A-B55A). Cells were harvested 48 h post-infection and lysed individually as described above. Lysates containing KBTBD4 were mixed 1:1 with PP2A-containing lysates or buffer respectively and were incubated for 1 h at 4°C. Complexes were captured using M2 resin (Sigma-Aldrich, A2220) to enrich for Flag-PP2A-A, StrepTactin-XT resin (IBA Lifesciences, 2-5010-025) to enrich for Strep-PP2A-C, or amylose resin (NEB, E8021L) to enrich for His-MBP-KBTBD4, followed by immunoprecipitation steps as described above.

### *In Vivo* Ubiquitylation

To detect specific ubiquitylation of PP2A-A in cells, Δ*KBTBD4* cells were transduced with lentivirus produced from pLVX-KBTBD4^3×Flag^-IRES-Puromycin constructs and selected with 1 μg mL^-1^ puromycin to express KBTBD4^WT^ or the cancer mutant KBTBD4^PR^. Control Δ*KBTBD4* cells and rescued cells were seeded at 4×10^6^ cells in 15 cm plates, and transiently transfected after 24 h with 9 μg of pCS2-^6×His^ubiquitin and 1.5 μg of pCS2-^myc^PP2A-A constructs using PEI (1:6 ratio). After 48 h of expression, cells were treated with 2 μM Carfilzomib (Cayman Chemicals, CAY17554) and 1 μM phorbol 12-myristate 13-acetate (PMA; Sigma-Aldrich, 79346) for 4 h, as indicated, harvested in cold PBS, and cell pellets were flash frozen. Samples were lysed in 1 mL of urea lysis buffer (8 M urea, 300 mM NaCl, 0.5% NP-40, 50 mM Na_2_HPO_4_, 50 mM Tris/HCl pH 8, 10 mM imidazole), supplemented with 10 mM *N*-ethylmaleimide (Sigma-Aldrich, 04259), Roche cOmplete protease inhibitor cocktail (Sigma-Aldrich, 5056489001) and phosphatase inhibitor cocktail (Thermo Fisher, A32957) and incubated at RT for 30 min. Samples were sonicated and centrifuged at 21,000×g for 10 min, and supernatants were normalized for protein concentration using Pierce 660 Protein Assay reagent (Thermo Fisher, 22660). For input, 5% of the supernatant was removed and the remaining sample was incubated with equilibrated Ni-NTA resin (Qiagen, 30230) at RT for 4 h. Resin was washed twice with lysis buffer and eluted with urea sample buffer supplemented with 200 mM imidazole. The eluates were subjected to SDS-PAGE and subsequent western blotting with the indicated antibodies.

### Sucrose Gradient Centrifugation

Cell extracts were prepared from 1.5×10^7^ cells of HEK293T and Δ*KBTBD4* cells by lysis in lysis buffer (20 mM HEPES pH7.4, 100 mM NaCl, 10 mM NaF, 1 mM Na_3_VO_4_, 2 mM phenanthroline, 0.2% NP40, protease inhibitor cocktail). Lysates were clarified by centrifugation for 20 min at 4,000×g. Clarified extracts were layered on top of a linear 5-20% (w/v) sucrose gradient prepared in gradient buffer (20 mM HEPES pH 7.4, 100 mM NaCl, 10 mM NaF, 1 mM sodium orthovanadate) and subjected to ultracentrifugation at 37,000rpm (Beckmann SW 60Ti rotor, 17h, 4°C). After centrifugation, 500 μL fractions were collected from the top of the gradient and fractions were analyzed by SDS-PAGE and western blotting with the indicated antibodies.

### Cellular Fitness Assay to Monitor Effects of Excess PP2A-A

To assess the effect of excess PP2A-A on cellular fitness, HEK293T cells were transduced with lentivirus encoding either pLVX-BFP-PP2A-A-IRES-mCherry-P2A-Blasticidin constructs (wild-type PP2A-A and the mutants P179R, R183W, S256F), or pLVX-GFP-P2A-Blasticidin control. Transduced cells were selected with 7.5 μg mL^-1^ blasticidin. After completion of selection, cells expressing BFP-PP2A-A-IRES-mCherry constructs were mixed with the GFP control population and passaged for 16 days under continuous blasticidin selection at 5 μg mL^-1^. At each passage, flow cytometry was used to measure the intracellular ratio of BFP/mCherry as a readout of PP2A-A protein stability, and the intercellular ratio of mCherry/GFP, as a measure of relative cellular fitness under PP2A-A overexpression conditions. Fitness measurements were normalized to the ratio of mCherry/GFP of day 0 for each sample. All experiments were performed in biological triplicates.

### Recombinant Protein Expression and Purification

Wild-type KBTBD4 and the indel cancer mutant PR, each carrying an N-terminal His-MBP tag followed by a TEV-cleavage site, were co-expressed with the chaperone NUDCD3 in insect cells. Baculoviruses encoding His-MBP-TEV-KBTBD4 and NUDCD3 were generated in Sf9 cells, and High Five cells were co-infected at a 1:1 ratio before harvesting 48 h later by centrifugation (1,000×g, 15 min). PP2A-A (WT or cancer mutants P179R or R183W) was expressed in *E. coli* BL21(DE3) Rosetta as GST-TEV-PP2A-A fusion. Cultures were grown at 37°C to OD₆₀₀≈0.6, induced with 0.6 mM IPTG, and expressed at 20°C for 16 h before harvesting (4,000×g, 15 min). Cell pellets were resuspended in lysis buffer (50 mM Tris-HCl pH 7.5, 200 mM NaCl, 0.5 mM TCEP, protease inhibitor cocktail) and lysed by sonication. Clarified lysates (50,000×g, 30 min, 4°C) were subjected to affinity purification using amylose resin (for KBTBD4; NEB, E8021L) or GSH resin (for PP2A-A; UBPBio, P3060). His-MBP-TEV-KBTBD4 was eluted with 10 mM maltose (Sigma-Aldrich, 63418) and cleaved overnight with TEV protease. TEV cleavage of PP2A-A was performed on resin overnight, and the cleaved protein was collected in the flow-through. Both, cleaved KBTBD4 and PP2A-A were further purified by anion-exchange chromatography (Cytiva, HiTrap Q HP) followed by size-exclusion chromatography (Cytiva, Superdex 200 10/300 GL) in 25 mM HEPES pH 7.5, 150 mM NaCl, 0.5 mM TCEP, aliquoted, flash frozen and stored at -80°C.

Purified PP2A-ACB55A, PP2A-ACB56E and PP2A-AC (sub-)complexes for ubiquitylation and mass photometry experiments were a generous gift from Franziska Wachter and Eric Fischer and prepared as described previously (Wachter et al. 2024).

For the E3 ligase complex, CUL3 with a C-terminal Strep-tag and untagged RBX1 were co-expressed in insect cells. Baculoviruses encoding CUL3-Strep or RBX1 were generated in Sf9 cells, and High Five cells were co-infected at a 3:1 ratio before harvesting 48 h later by centrifugation (1000×g, 15 min). Cell pellets were processed as described above. Clarified lysate was subjected to affinity purification using Strep-Tactin-resin (IBA Lifesciences, 2-1201-010). The CUL3-Strep-RBX1 complex was eluted with 6 mM desthiobiotin (IBA Lifesciences, 2-1000-002) and further purified by cation-exchange chromatography (Cytiva, HiTrap SP HP). A final concentration of 6 μM of the CUL3-Strep-RBX1 complex was neddylated with 30 μM commercially available NEDD8 (R&D Systems, UL-812) by 0.2 μM UBA3 (R&D Systems, E-313-025) and 1.2 μM UBE2M (R&D Systems, E2-656-100) after addition of 2.5 mM ATP and 2.5 mM MgCl_2_ in 50 mM Tris-HCl pH 8 and 150 mM NaCl for 30 min at RT. The neddylated CUL3-Strep-RBX1 complex was purified by size-exclusion chromatography (Cytiva, Superdex 200 10/300 GL) in 25 mM HEPES pH 7.5, 150 mM NaCl, 0.5 mM TCEP, aliquoted, flash frozen and stored at -80°C.

### *In vitro* Ubiquitylation

Ubiquitylation assays were carried out in 20 μL reactions with final concentrations of 1 μM PP2A (PP2A-A or in indicated complex), 0.1 μM UBE1 (R&D Systems, E-304), 0.2 μM UBE2D3 (R&D Systems, E2-627), 0.5 μM KBTBD4, 0.4 μM CUL3^NEDD8^-RBX1 (CRL), 7.5 μM ubiquitin^IRdye680^ (gift from Jake Aguirre, as published in (Tsai et al. 2023)), 5 mM ATP, 2.5 mM MgCl_2_, in a buffer of 20 mM HEPES pH 7.4, 150 mM NaCl and 1 mM DTT.

Reactions were carried out at RT for 30 min and quenched by addition of 20 μL 2×USB + 5% BME. The samples were analyzed by SDS-PAGE and fluorescent signal of ubiquitylated species was subsequently imaged with an Odyssey CLx Imager (LI-COR Biotech). The assay was additionally analyzed by western blot for PP2A-A, as well as coomassie staining for total protein loading.

### Mass Photometry

KBTBD4 and PP2A-A or PP2A (sub-)complexes were diluted in MP buffer (25 mM HEPES, 150 mM NaCl, 0.5 mM TCEP) to concentrations of 0.5 μM in 10 μL, or mixed at a 1:1 ratio at 0.5 μM each in 10 μL and incubated on ice for 15 min. For data acquisition on a Refeyn OneMP mass photometer, 19 μL of MP buffer was added to the flow chamber, followed by focus calibration. Next, 1 μL of the protein solution was added to the chamber to acquire movies of 60 s at a final concentration of 25 nM. Each sample was measured at least two times, and acquired movies were processed, and molecular masses were analyzed using Refeyn Discoverer 2.3 software based on the BSA standard curve.

### Cryo Electron Microscopy

To determine the binding mode of KBTBD4 to its endogenous substrate PP2A-A, the complex was analyzed by cryo-EM. KBTBD4 and PP2A-A (each at 0.6 mg mL⁻¹) were cross-linked with BS3, and the reaction was quenched after 30 min by addition of 50 mM Tris-HCl pH 7.5. Immediately prior to vitrification, n-octyl-β-D-glucopyranoside (β-OG) was added to a final concentration of 0.1%. 3.5 μL sample was applied to freshly glow-discharged Quantifoil R1.2/1.3 Cu 300 grids using a Vitrobot Mark IV (Thermo Fisher Scientific) at 100% humidity and 4°C, blotted for 4 s at blot force 4, and plunge-frozen in liquid ethane. Data were collected on a 300kV Titan Krios G4 transmission electron microscope (Thermo Fisher Scientific) equipped with a E-CFEG (low energy spread cold field emission gun), Falcon 4i direct electron detector and a SelectrisX energy filter (10eV), yielding 20,258 micrographs at a nominal magnification of 165,000, pixel size of 0.732 Å, defocus range of 0.8-2.2 μm, and total exposure of 40e⁻ per Å^2^.

All data processing was performed in CryoSPARC v4.7 (Punjani et al. 2017) unless noted otherwise. Other software was compiled and configured using SBGrid (Morin et al. 2013). Following motion correction and contrast transfer function (CTF) estimation, 19,923 micrographs were retained. An *ab initio* model was generated from a subset of 1,000 micrographs using the Blob Picker and 2D classification. This map served as the initial model for further processing. Using the full dataset, 983,288 particles were picked with the Blob Picker and extracted, followed by 2D classification, several iterations of non-uniform, and homogeneous refinements, and local and global CTF refinement (Punjani et al. 2020) (**Fig. S3**). The final reconstruction of a KBTBD4 dimer bound to two PP2A-A monomers was obtained at 2.62 Å resolution after application of C2 symmetry. DeepEMhancer was used to improve map features for visualization (Sanchez-Garcia et al., 2021). Heterogeneous refinement revealed conformational heterogeneity between the two PP2A-A moieties and a focused map was obtained after heterogeneous and local refinement to improve the edges of the map. After extensive heterogeneous, non-uniform, and homogeneous refinement, a second reconstruction corresponding to a KBTBD4 dimer bound to a single PP2A-A monomer was obtained at 3.7 Å resolution.

For model building, an AlphaFold3 model (Abramson et al. 2024) of a single PP2A-A with a KBTBD4 dimer was used as a starting point. Model building was done in ChimeraX (Meng et al. 2023) using the ISOLDE plugin (Croll 2018) on the main map from CryoSPARC and the focused map on the edges of the model where loops were not well resolved. This was followed by several rounds of refinement using Phenix (real_space_refine) (Liebschner et al. 2019; Afonine et al. 2018) and manual inspection in ISOLDE.

### RQ-TRAP Telomerase Activity Assay

Relative telomerase activity was measured in biological triplicates by RQ-TRAP, a modified version of the TRAP assay, in which telomere elongation products are amplified by quantitative RT-PCR (Wege et al. 2003). HEK293T and Δ*KBTBD4* cells were harvested in PBS, flash frozen in liquid nitrogen and stored at -80°C. Pellets were resuspended in CHAPS lysis buffer (10 mM Tris-HCl pH 7.5, 1 mM MgCl_2_, 1 mM EGTA, 5 mM BME, 0.5% CHAPS, 10% glycerol) at 1×10^6^ cells per mL. After 30 min incubation on a rotating wheel at 4°C, extracts were centrifuged at 14,000×g for 30 min at 4°C, the supernatants were flash frozen in liquid nitrogen and stored at -80°C. Protein concentration of extracts was determined with the Bradford protein assay (BioRad).

Telomerase activity reactions were carried out in 20 μL volumes containing 10 μL of Applied Biosystems Power SYBR Green PCR master mix (Thermo Fisher, 4367659), 2 μL of 400 ng μL^-1^ cell extracts (or serial dilution), 0.1 μg of telomerase primer TS (5’-AAT CCG TCG AGC AGA GTT-3’), 0.1 μg of reverse primer ACX (5’-GCG CGG CTT ACC CTT ACC CTT ACC CTA ACC-3’) and 2 μL of 10 mM EGTA. Samples were incubated for 30 min at 30°C protected from light for telomerase extension: 10 min at 95°C and telomerase products were amplified by 40 PCR cycles consisting of 15 s at 95°C and 60 s at 60°C using the QuantStudio 7 instrument (Thermo Fisher). The threshold cycle values (*C*_t_) were determined in triplicate for each sample. Duplicate five-fold serial dilutions of the WT reference sample were used to draw a standard curve of the form log_10_(protein quantity)=*aC*_t_+*b*. Telomerase activity was expressed relative to this standard as the quantity of standard sample extract giving the same *C*_t_ value. Samples were serially diluted to verify the linearity of the RQ-TRAP reaction, and RNase-treated and/or heat inactivated to verify that the amplification products were due to telomerase activity.

### Telomere Restriction Fragment Length Analysis

Genomic DNA from 3-5×10^6^ cells was extracted with phenol-chloroform-isoamyl alcohol (25:24:1) and precipitated with isopropanol. 5-10 μg of genomic DNA were digested with 50 U HinfI and 30 U RsaI in CutSmart buffer (NEB) overnight at 37°C. 3 μg of digested genomic DNA per lane was loaded on a 0.8% agarose gel in 1×TBE and run at 2.4 v/cm for 16 h. The gel was stained with gel red for 40 min to detect total DNA and size markers. The gel was quickly rinsed and dried for 3 h at 50°C on a BioRad gel dryer. The dried gel was denatured in 0.5 M NaOH, 1.5 M NaCl for 15 min and neutralized in 0.5 M Tris-HCl pH 7.5, 1.5 M NaCl for 15 min. The gel was then prehybridized with Church buffer (0.5 M Na_2_HPO_4_ pH 7.2, 1 mM EDTA, 7% SDS, and 1% BSA) for 1 h at 50°C and hybridized with a ^32^P-radiolabeled telomeric probe overnight at 50°C. After hybridization, the gel was washed at 50°C in 4× SSC for 1 h, once in 4× SSC /0.1% SDS for 1 h, once in 2× SSC/0.1% SDS for 1 h, and then quickly rinsed with 4× SSC. The washed gel was exposed, and radioactive signals were detected with an Amersham typhoon phosphorimager.

### Resource Availability

The atomic coordinates have been deposited in the Protein Data Bank with accession code PDB: 28JO. The cryo-EM maps included in this study have been deposited in the Electron Microscopy Data Bank under the accession codes EMDB: EMD-56545 (main map for 2:2 complex), EMD-54547 (main map for 2:1 complex), and EMD-56548 (focused map for 2:2 complex). The PDB and EMDB entries will be openly available upon publication, and the maps will be made available to reviewers upon request.

The proteomics and phospho proteomics data will be deposited in the Proteomics Identification Database (PRIDE).

This paper does not report original code.

## Acknowledgements

We are grateful to all Rape and Thomä lab members for helpful discussions, and to Achim Werner, Frauke Melchior, and Julian Yano for insights and suggestions. We thank Franziska Wachter and Eric Fischer for purified PP2A complexes and helpful discussions. We thank Fiona Bello and Mira Schütz for laboratory management and organization. Cryo-EM data collection was performed at the Dubochet Center for Imaging Lausanne (a common initiative from EPFL, UNIGE, UNIL, UNIBE). Data processing was performed using SCITAS infrastructure at EPFL. We gratefully acknowledge A. Atrih, Filipe Soares R. from the Dundee FingerPrints Proteomics Facility for their excellent technical support.

## Funding

N.H.T.’s laboratory was supported by funding from the European Research Council (ERC) under the European Union’s H2020 research program (NucEM; 884331), the Swiss National Science Foundation (SNSF; 310030_301206 and 310030_214852), Krebsforschung (KFS; KFS-5933-08-2023), and the Mark Foundation (Aspire Award 10911). M.R. is an Investigator of the Howard Hughes Medical Institute. J.L.’s laboratory was supported by the SNSF (310030_214833). A.C.’s laboratory has received funding from the Innovative Medicines Initiative 2 (IMI2) Joint Undertaking under Grant 875510 (EUbOPEN project). The IMI2 Joint Undertaking receives support from the European Union’s Horizon 2020 research and innovation program, EFPIA companies, and associated partners: KTH, OICR, Diamond, and McGill.

## Conflict of interest

The Thomä laboratory receives industrial funding from the Novartis Research Foundation, AstraZeneca, Merck KGaA and Sanofi-Aventis. N.H.T. has consulted for Monte Rosa, Boehringer Ingelheim, Astra Zeneca, Ridgeline Therapeutics, Red Ridge Bio, and is a founder and shareholder of Zenith Therapeutics. M.R. is cofounder and SAB member of Nurix Therapeutics; co-founder, consultant and SAB chair for Lyterian Therapeutics, co-founder and consultant of Zenith Therapeutics, co-founder of Reina, and iPartner of The Column Group. The Ciulli laboratory receives or has received sponsored research support from Almirall, Amgen, Amphista Therapeutics, Boehringer Ingelheim, Eisai, Merck KGaA, Nurix Therapeutics, Ono Pharmaceuticals and Tocris-Biotechne. A.C. is a scientific founder and shareholder of Amphista Therapeutics, a company that is developing targeted protein degradation therapeutic platforms, and is on the Scientific Advisory Board of ProtOS Bio and TRIMTECH Therapeutics.

## References

Abramson, Josh, Jonas Adler, Jack Dunger, et al. 2024. “Accurate Structure Prediction of Biomolecular Interactions with AlphaFold 3.” Nature 630 (8016): 493–500. 10.1038/s41586-024-07487-w.

Afonine, P. V., B. K. Poon, R. J. Read, et al. 2018. “Real-Space Refinement in PHENIX for Cryo-EM and Crystallography.” Acta Crystallographica Section D: Structural Biology 74 (6): 531–44. 10.1107/S2059798318006551.

Akopian, David, Colleen A McGourty, and Michael Rape. 2022. Co-Adaptor Driven Assembly of a CUL3 E3 Ligase Complex Ll Ll Co-Adaptor Driven Assembly of a CUL3 E3 Ligase Complex. 585–97. 10.1016/j.molcel.2022.01.004.

Andrés, María E., Corinna Burger, María J. Peral-Rubio, et al. 1999. “CoREST: A Functional Corepressor Required for Regulation of Neural-Specific Gene Expression.” Proceedings of the National Academy of Sciences 96 (17): 9873–78. 10.1073/pnas.96.17.9873.

Brautigan, David L., and Shirish Shenolikar. 2018. “Protein Serine/Threonine Phosphatases: Keys to Unlocking Regulators and Substrates.” Annual Review of Biochemistry 87 (1): 921–64. 10.1146/annurev-biochem-062917-012332.

Brewer, Abigail, Gajanan Sathe, B Pflug, R Clarke, T Macartney, and G Sapkota. 2024. “Mapping the Substrate Landscape of Protein Phosphatase 2A Catalytic Subunit PPP2CA.” iScience 27 (3). 10.1016/j.isci.2024.109302.

Cavalli, Florence M.G., Marc Remke, Ladislav Rampasek, et al. 2017. “Intertumoral Heterogeneity within Medulloblastoma Subgroups.” Cancer Cell 31 (6): 737–754.e6. 10.1016/j.ccell.2017.05.005.

Chagraoui, Jalila, Simon Girard, Jean-Francois Spinella, et al. 2021. “UM171 Preserves Epigenetic Marks That Are Reduced in Ex Vivo Culture of Human HSCs via Potentiation of the CLR3-KBTBD4 Complex.” Cell Stem Cell 28 (1): 48–62.e6. 10.1016/j.stem.2020.12.002.

Chen, Zhuoyao, Gamma Chi, Timea Balo, et al. 2025. “Structural Mimicry of UM171 and Neomorphic Cancer Mutants Co-Opts E3 Ligase KBTBD4 for HDAC1/2 Recruitment.” Nature Communications 16 (1): 3144. 10.1038/s41467-025-58350-z.

Chen, Zhuoyao, Rafael M. Ioris, Stacey Richardson, et al. 2022. “Disease-Associated KBTBD4 Mutations in Medulloblastoma Elicit Neomorphic Ubiquitylation Activity to Promote CoREST Degradation.” Cell Death & Differentiation 29 (10): 10. 10.1038/s41418-022-00983-4.

Cho, Uhn Soo, and Wenqing Xu. 2007. “Crystal Structure of a Protein Phosphatase 2A Heterotrimeric Holoenzyme.” Nature 445 (7123): 7123. 10.1038/nature05351.

Cong, Le, F. Ann Ran, David Cox, et al. 2013. “Multiplex Genome Engineering Using CRISPR/Cas Systems.” Science 339 (6121): 819–23. 10.1126/science.1231143.

Croll, T. I. 2018. “ISOLDE: A Physically Realistic Environment for Model Building into Low-Resolution Electron-Density Maps.” Acta Crystallographica Section D: Structural Biology 74 (6): 519–30. 10.1107/S2059798318002425.

Curtin, Sally C, Minino, Arialdi M, and Anderson, Robert N. 2016. “Declines in Cancer Death Rates Among Children and Adolescents in the United States, 1999–2014.” NCHS Data Brief, no. 257.

Fowle, Holly, Ziran Zhao, and Xavier Graña. 2019. “PP2A Holoenzymes, Substrate Specificity Driving Cellular Functions and Deregulation in Cancer.” Advances in Cancer Research 144: 55–93. 10.1016/bs.acr.2019.03.009.

Groves, Matthew R., Neil Hanlon, Patric Turowski, Brian A. Hemmings, and David Barford. 1999. “The Structure of the Protein Phosphatase 2A PR65/A Subunit Reveals the Conformation of Its 15 Tandemly Repeated HEAT Motifs.” Cell 96 (1): 99–110. 10.1016/S0092-8674(00)80963-0.

Haakonsen, Diane L., Michael Heider, Andrew J. Ingersoll, et al. 2024. “Stress Response Silencing by an E3 Ligase Mutated in Neurodegeneration.” *Nature*, January 31, 1– 7. 10.1038/s41586-023-06985-7.

Haanen, Terrance J., Caitlin M. O’Connor, and Goutham Narla. 2022. “Biased Holoenzyme Assembly of Protein Phosphatase 2A (PP2A): From Cancer to Small Molecules.” The Journal of Biological Chemistry 298 (12): 102656. 10.1016/j.jbc.2022.102656.

Ingersoll, Andrew J., Devlon M. McCloud, Jenny Y. Hu, and Michael Rape. 2025. “Dynamic Regulation of the Oxidative Stress Response by the E3 Ligase TRIP12.” Cell Reports 44 (9): 116262. 10.1016/j.celrep.2025.116262.

Ismail, Houssam, Jalila Chagraoui, and Guy Sauvageau. 2025. “CoREST in Pieces: Dismantling the CoREST Complex for Cancer Therapy and Beyond.” Science Advances 11 (23): eads6556. 10.1126/sciadv.ads6556.

Kauko, Otto, and Jukka Westermarck. 2018. “Non-Genomic Mechanisms of Protein Phosphatase 2A (PP2A) Regulation in Cancer.” International Journal of Biochemistry and Cell Biology 96 (December 2017): 157–64. 10.1016/j.biocel.2018.01.005.

Kong, Mei, Dara Ditsworth, Tullia Lindsten, and Craig B. Thompson. 2009. “Α4 Is an Essential Regulator of PP2A Phosphatase Activity.” Molecular Cell 36 (1): 51–60. 10.1016/j.molcel.2009.09.025.

Lee, Julieann C., Tali Mazor, Richard Lao, et al. 2019. “Recurrent KBTBD4 Small In-Frame Insertions and Absence of DROSHA Deletion or DICER1 Mutation Differentiate Pineal Parenchymal Tumor of Intermediate Differentiation (PPTID) from Pineoblastoma.” Acta Neuropathologica 137 (5): 851–54. 10.1007/s00401-019-01990-5.

Li, H., L. L. Zhao, J. W. Funder, and J. P. Liu. 1997. “Protein Phosphatase 2A Inhibits Nuclear Telomerase Activity in Human Breast Cancer Cells.” The Journal of Biological Chemistry 272 (27): 16729–32. 10.1074/jbc.272.27.16729.

Li, Mei, Anthony Makkinje, and Zahi Damuni. 1996. “The Myeloid Leukemia-Associated Protein SET Is a Potent Inhibitor of Protein Phosphatase 2A (∗).” Journal of Biological Chemistry 271 (19): 11059–62. 10.1074/jbc.271.19.11059.

Liebschner, D., P. V. Afonine, M. L. Baker, et al. 2019. “Macromolecular Structure Determination Using X-Rays, Neutrons and Electrons: Recent Developments in Phenix.” Acta Crystallographica Section D: Structural Biology 75 (10): 861–77. 10.1107/S2059798319011471.

Liu, Chunming, Yoichi Kato, Zhuohua Zhang, Viet Minh Do, Bruce A. Yankner, and Xi He. 1999. “β-Trcp Couples β-Catenin Phosphorylation-Degradation and Regulates Xenopus Axis Formation.” Proceedings of the National Academy of Sciences 96 (11): 6273–78. 10.1073/pnas.96.11.6273.

Maiwald, Samuel A., Laura A. Schneider, Ronnald Vollrath, et al. 2025. “TRIP12 Structures Reveal HECT E3 Formation of K29 Linkages and Branched Ubiquitin Chains.” Nature Structural & Molecular Biology 32 (9): 1766–75. 10.1038/s41594-025-01561-1.

Manford, Andrew G., Fernando Rodríguez-Pérez, Karen Y. Shih, et al. 2020. “A Cellular Mechanism to Detect and Alleviate Reductive Stress.” Cell 183 (1): 46–61.e21. 10.1016/j.cell.2020.08.034.

Mark, Kevin G., SriDurgaDevi Kolla, Jacob D. Aguirre, et al. 2023. “Orphan Quality Control Shapes Network Dynamics and Gene Expression.” *Cell*, July, S0092867423006918. 10.1016/j.cell.2023.06.015.

Mena, Elijah L., Rachel A. S. Kjolby, Robert A. Saxton, et al. 2018. “Dimerization Quality Control Ensures Neuronal Development and Survival.” Science 362 (6411): eaap8236. 10.1126/science.aap8236.

Meng, Elaine C., Thomas D. Goddard, Eric F. Pettersen, et al. 2023. “UCSF ChimeraX: Tools for Structure Building and Analysis.” Protein Science 32 (11): e4792. 10.1002/pro.4792.

Morin, Andrew, Ben Eisenbraun, Jason Key, et al. 2013. “Collaboration Gets the Most out of Software.” eLife 2 (September): e01456. 10.7554/eLife.01456.

Northcott, Paul A., Ivo Buchhalter, A. Sorana Morrissy, et al. 2017. “The Whole-Genome Landscape of Medulloblastoma Subtypes.” Nature 547 (7663): 7663. 10.1038/nature22973.

Northcott, Paul A., Adrian M. Dubuc, Stefan Pfister, and Michael D. Taylor. 2012. “Molecular Subgroups of Medulloblastoma.” Expert Review of Neurotherapeutics 12 (7): 871–84. 10.1586/ern.12.66.

Northcott, Paul A., Andrey Korshunov, Hendrik Witt, et al. 2011. “Medulloblastoma Comprises Four Distinct Molecular Variants.” Journal of Clinical Oncology 29 (11): 1408–14. 10.1200/JCO.2009.27.4324.

Northcott, Paul A., Catherine Lee, Thomas Zichner, et al. 2014. “Enhancer Hijacking Activates GFI1 Family Oncogenes in Medulloblastoma.” Nature 511 (7510): 428–34. 10.1038/nature13379.

Northcott, Paul A., David J. H. Shih, John Peacock, et al. 2012. “Subgroup-Specific Structural Variation across 1,000 Medulloblastoma Genomes.” Nature 488 (7409): 7409. 10.1038/nature11327.

O’Connor, Caitlin M., Daniel Leonard, Danica Wiredja, et al. 2020. “Inactivation of PP2A by a Recurrent Mutation Drives Resistance to MEK Inhibitors.” Oncogene 39 (3): 703–17. 10.1038/s41388-019-1012-2.

Padovani, Chris, Predrag Jevtić, and Michael Rapé. 2022. “Quality Control of Protein Complex Composition.” Molecular Cell 82 (8): 1439–50. 10.1016/j.molcel.2022.02.029.

Park, Jisu, Kyubin Lee, Kyunghwan Kim, and Sun-Ju Yi. 2022. “The Role of Histone Modifications: From Neurodevelopment to Neurodiseases.” Signal Transduction and Targeted Therapy 7 (1): 217. 10.1038/s41392-022-01078-9.

Pavic, Karolina, Nikhil Gupta, Judit Domènech Omella, et al. 2023. “Structural Mechanism for Inhibition of PP2A-B56α and Oncogenicity by CIP2A.” Nature Communications 14 (1): 1143. 10.1038/s41467-023-36693-9.

Punjani, Ali, John L. Rubinstein, David J. Fleet, and Marcus A. Brubaker. 2017. “cryoSPARC: Algorithms for Rapid Unsupervised Cryo-EM Structure Determination.” Nature Methods 14 (3): 290–96. 10.1038/nmeth.4169.

Punjani, Ali, Haowei Zhang, and David J. Fleet. 2020. “Non-Uniform Refinement: Adaptive Regularization Improves Single-Particle Cryo-EM Reconstruction.” Nature Methods 17 (12): 1214–21. 10.1038/s41592-020-00990-8.

Rodríguez Pérez, Fernando, Dean Natwick, Lauren Schiff, et al. 2024. “WRN Inhibition Leads to Its Chromatin-Associated Degradation via the PIAS4-RNF4-P97/VCP Axis.” Nature Communications 15 (1): 6059. 10.1038/s41467-024-50178-3.

Rodríguez-Pérez, Fernando, Andrew G. Manford, Angela Pogson, Andrew J. Ingersoll, Brenda Martínez-González, and Michael Rape. 2021. “Ubiquitin-Dependent Remodeling of the Actin Cytoskeleton Drives Cell Fusion.” Developmental Cell 56 (5): 588–601.e9. 10.1016/j.devcel.2021.01.016.

Sangodkar, Jaya, Caroline C. Farrington, Kimberly McClinch, Matthew D. Galsky, David B. Kastrinsky, and Goutham Narla. 2016. “All Roads Lead to PP2A: Exploiting the Therapeutic Potential of This Phosphatase.” FEBS Journal 283 (6): 1004–24. 10.1111/febs.13573.

Shannon, Paul, Andrew Markiel, Owen Ozier, et al. 2003. “Cytoscape: A Software Environment for Integrated Models of Biomolecular Interaction Networks.” Genome Research 13 (11): 2498–504. 10.1101/gr.1239303.

Sharma, Tanvi, Edward C. Schwalbe, Daniel Williamson, et al. 2019. “Second-Generation Molecular Subgrouping of Medulloblastoma: An International Meta-Analysis of Group 3 and Group 4 Subtypes.” Acta Neuropathologica 138 (2): 309–26. 10.1007/s00401-019-02020-0.

Taylor, Sarah E., Caitlin M O Connor, Zhizhi Wang, et al. 2019. “The Highly Recurrent PP2A Aa-Subunit Mutation P179R Alters Protein Structure and Impairs PP2A Enzyme Function to Promote Endometrial Tumorigenesis.” Cancer Research 79 (16): 4242–57. 10.1158/0008-5472.CAN-19-0218.

Tsai, Jonathan M., Jacob D. Aguirre, Yen-Der Li, et al. 2023. “UBR5 Forms Ligand-Dependent Complexes on Chromatin to Regulate Nuclear Hormone Receptor Stability.” Molecular Cell 0 (0). 10.1016/j.molcel.2023.06.028.

Wachter, Franziska, Radosław P. Nowak, Scott Ficarro, Jarrod Marto, and Eric S. Fischer. 2024. “Structural Characterization of Methylation-Independent PP2A Assembly Guides alphafold2Multimer Prediction of Family-Wide PP2A Complexes.” Journal of Biological Chemistry 300 (5): 107268. 10.1016/j.jbc.2024.107268.

Wege, Henning, Michael S. Chui, Hai T. Le, Julie M. Tran, and Mark A. Zern. 2003. “SYBR Green Real-time Telomeric Repeat Amplification Protocol for the Rapid Quantification of Telomerase Activity.” Nucleic Acids Research 31 (2): e3. 10.1093/nar/gng003.

Werner, Achim, Regina Baur, Nia Teerikorpi, Deniz U. Kaya, and Michael Rape. 2018. “Multisite Dependency of an E3 Ligase Controls Monoubiquitylation-Dependent Cell Fate Decisions.” eLife 7: 1–27. 10.7554/eLife.35407.

Wu, Chang-Chih, Shirui Hou, Brent A. Orr, et al. 2017. “mTORC1-Mediated Inhibition of 4EBP1 Is Essential for Hedgehog Signaling-Driven Translation and Medulloblastoma.” Developmental Cell 43 (6): 673–688.e5. 10.1016/j.devcel.2017.10.011.

Xie, Xiaowen, Olivia Zhang, Megan J. R. Yeo, et al. 2025. “Converging Mechanism of UM171 and KBTBD4 Neomorphic Cancer Mutations.” Nature 639 (8053): 241–49. 10.1038/s41586-024-08533-3.

Xu, Yanhui, Yongna Xing, Yu Chen, et al. 2006. “Structure of the Protein Phosphatase 2A Holoenzyme.” Cell 127 (6): 1239–51. 10.1016/j.cell.2006.11.033.

Yen, Hsueh-Chi Sherry, and Stephen J. Elledge. 2008. “Identification of SCF Ubiquitin Ligase Substrates by Global Protein Stability Profiling.” Science 322 (5903): 923–29. 10.1126/science.1160462.

Yen, Hsueh-Chi Sherry, Qikai Xu, Danny M. Chou, Zhenming Zhao, and Stephen J. Elledge. 2008. “Global Protein Stability Profiling in Mammalian Cells.” *Science (New York*, N.Y.) 322 (5903): 918–23. 10.1126/science.1160489.

Yeo, Megan J. R., Olivia Zhang, Xiaowen Xie, et al. 2025. “UM171 Glues Asymmetric CRL3–HDAC1/2 Assembly to Degrade CoREST Corepressors.” Nature 639 (8053): 232–40. 10.1038/s41586-024-08532-4.

